# HIF2-driven PTHrP Causes Cachexia and Hypercalcemia in Kidney Cancer: Treatment with HIF2 Inhibitors

**DOI:** 10.1101/2025.09.09.675147

**Authors:** Muhannad Abu-Remaileh, Laura A. Stransky, Nikita Bhalerao, Nitin H. Shirole, Qinqin Jiang, Eddy Saad, Marc Machaalani, Sean M. Vigeant, Hilina Woldemichael, Charles Xu, Jing Lu, Hairong Wei, Zhihong Liu, William Sun, Kei Enomoto, Toni K. Choueiri, Jason R. Pitarresi, Steven A. Carr, Namrata D. Udeshi, William G. Kaelin

**Affiliations:** Department of Medical Oncology, Dana-Farber Cancer Institute, Harvard Medical School, Boston, MA 02215, USA; Broad Institute of MIT and Harvard, 415 Main Street, Cambridge, MA 02142, USA; Division of Hematology/Oncology, Department of Medicine, University of Massachusetts Chan Medical School, Worcester, MA, USA; Department of Medicine, Beth Israel Deaconess Medical Center, Harvard Medical School, Boston, MA, USA; NiKang Therapeutics 200 Powder Mill Rd E500, Wilmington, DE 19803, USA; Howard Hughes Medical Institute, Chevy Chase, MD 20815, USA

## Abstract

Kidney cancer frequently causes paraneoplastic syndromes, including hypercalcemia and cachexia, but the underlying mechanisms are incompletely understood. The most common form of kidney cancer, clear cell renal cell carcinoma, is frequently caused by loss of the pVHL tumor suppressor protein and the resulting upregulation of the HIF2 transcription factor. We show that *PTHLH*, which resides on a ccRCC amplicon on chromosome 12p, is a direct HIF2 transcriptional target in ccRCC. Further, we show that the increased *PTHLH* expression is both necessary and sufficient for the induction of hypercalcemia and cachexia in preclinical orthotopic cell line tumor models. Consistent with these observations, two different allosteric HIF2 inhibitors, belzutifan and NKT2152, rapidly ameliorated hypercalcemia and cachexia in ccRCC patients, including in some patients who did not exhibit objective tumor shrinkage.

## Introduction

Kidney cancer is one of the ten most common cancers in the United States, with ∼81,500 new cases and 14,000 deaths each year ^1^. Worldwide, an estimated 175,000 people died of kidney cancer in 2020 ^2^. Although early stage kidney cancer can often be cured surgically, treating advanced stage kidney cancer and its complications remains a challenge ^1^.

By far the most common form of kidney cancer is clear cell renal cell carcinoma (ccRCC) ^3^. The initiating molecular event in most ccRCCs is biallelic inactivation of the von Hippel-Lindau (*VHL*) tumor suppressor gene resulting from *VHL* mutations or *VHL* hypermethylation ^4^. The *VHL* gene product, pVHL, is the substrate recognition subunit of an E3 ubiquitin ligase that targets the alpha subunits of the heterodimeric transcription factor HIF (Hypoxia-Inducible Factor) for proteasomal degradation when oxygen is abundant. In *VHL* mutant ccRCCs, loss of pVHL function causes the inappropriate accumulation of HIF and HIF target genes even under normoxic conditions ^5^. In preclinical models, HIF2, the transcription factor formed by HIF2α and an ARNT (Aryl Hydrocarbon Receptor Nuclear Translocator) protein, and not the canonical HIF family member HIF1α, drives ccRCC proliferation *ex vivo* and *in vivo* ^5,6^. Drugs that block the HIF2-responsive growth factor VEGF (vascular endothelial growth factor) or its receptor KDR (kinase-domain related) are now mainstays of ccRCC treatment ^7^. Upregulation of HIF2-responsive endogenous retroviruses might also contribute to the responsiveness of ccRCCs to immunotherapy ^8^. Allosteric HIF2 inhibitors were also recently shown to be active against a subset of heavily pretreated ccRCC patients ^9^. The first of these, belzutifan (Welireg), is now approved for the treatment of sporadic ccRCCs and tumor arising in patients with germline *VHL* mutations (VHL Disease) ^10,11^.

Kidney cancer is sometimes called “The Internist’s Tumor” since some kidney cancer patients first come to medical attention because they suffer from one of its many paraneoplastic complications. Examples of such complications include fever of unknown origin, elevated erythrocyte sedimentation rate, hypertension, hypercalcemia, anemia, polycythemia, abnormal liver function tests without metastases (remote hepatopathy or Stauffer’s Syndrome), coagulopathy, altered glucose metabolism, galactorrhea, amyloidosis, and Cushing’s Syndrome ^12–14^. Polycythemia in the setting of ccRCC is almost certainly caused by ectopic production of the HIF2-responsive growth factor erythropoietin ^15^. The molecular basis for most of the remaining ccRCC paraneoplastic syndromes remains obscure.

Cachexia, a multifactorial systemic syndrome that involves crosstalk between multiple organ systems across the body and characterized by involuntary loss of muscle mass and fat leading to >5% body weight reduction, is a common complication of chronic conditions such as heart failure, chronic obstructive pulmonary disease, and cancer ^16^. This syndrome significantly diminishes quality of life, impairs tolerance to anticancer therapies, and accounts for up to 20% of cancer-related deaths ^17^. Notably, up to 35% of renal cell carcinoma patients develop cachexia ^13,14^.

PTHrP (parathyroid hormone-related protein) has been implicated in cachexia caused by some cancers and by kidney failure ^18–23^. Here we show that the PTHrP, which is encoded by the HIF2-responsive gene *PTHLH*, is a critical driver of two paraneoplastic syndromes in ccRCC: cachexia and humoral hypercalcemia. Using preclinical and clinical models, we demonstrate that both syndromes can be ameliorated with clinical grade HIF2 inhibitors, underscoring the therapeutic potential of targeting HIF2-driven pathways in ccRCC-associated paraneoplastic syndromes.

## Results

HIF2 inhibitors inhibit tumor growth and prolong survival in multiple preclinical ccRCC models, including both orthotopic cell line models and PDX models ^24,25^. The HIF2 inhibitor PT2399 is a tool compound that is closely related to the approved drug belzutifan. We reported that PT2399 dramatically improved the survival of female nude mice bearing orthotopic tumors formed by the OSRC-2 *VHL-/-* ccRCC cell line (Extended Data Fig. 1a) ^26^. In this setting, however, PT2399 did not profoundly suppress tumor growth (Extended Data Fig. 1b). This was surprising because OSRC-2 cells are highly HIF-2-dependent *in vitro* based on 2D and 3D cell culture proliferation assays ^26^. Interestingly, we noticed that mice bearing OSRC-2 developed profound cachexia (Extended Data Fig. 1c), which ultimately required that they be euthanized, unless they were treated with PT2399.

To study this observation prospectively, we performed additional OSRC-2 subcutaneous xenograft assays, monitoring body weight weekly. As expected, the tumor-bearing mice developed profound cachexia. Once the mice lost more than 10% of their body weights, they were randomized to PT2399 or vehicle by oral gavage (Fig. 1a). Mice treated with PT2399 rapidly regained weight, while the vehicle-treated mice continued to lose weight (Fig. 1b,c and Extended Data Fig. 1d). We weighed tumors removed at necropsies after 6 days of therapy or, in the case of one vehicle-treated mouse, after 5 days to prevent animal suffering. The PT2399-treated tumors were not smaller, and trended to be larger, than the vehicle-treated tumors, suggesting that the acute reversal of cachexia by PT2399 was pharmacodynamic and not due to a loss of tumor cells (Fig. 1d). Immunoblot assays confirmed other pharmacodynamic (PD) effects of PT2399, including downregulation of Cyclin D1, NDRG1, and HIF2 protein levels itself ^25–27^ (Fig. 1e). The latter has been described before and likely reflects destabilization of monomeric HIF2α when it is no longer bound to ARNT.

**Figure 1.**
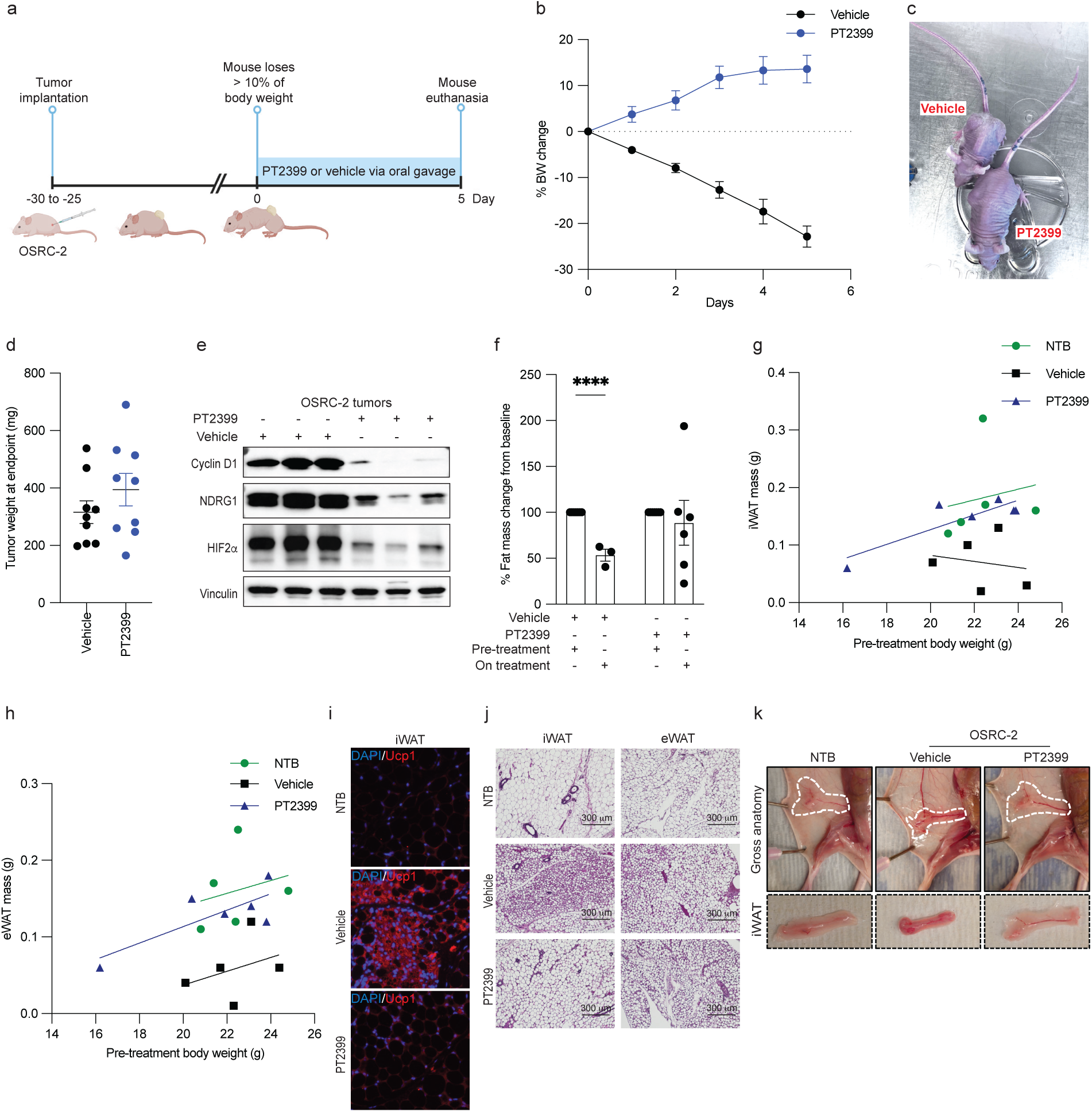
Successful Treatment of Cachexia Caused by OSRC-2 Xenografts with a HIF2 Inhibitor. (a) Treatment schema. (b) Body weight (BW) changes over time in OSRC-2 tumor-bearing mice treated with PT2399 (blue, n = 9) or vehicle (black, n = 9). (c and d) Representative mouse images (c) and tumor weights (d) at study endpoint of mice from (b). (e) Immunoblot analysis of representative OSRC-2 tumor lysates at study endpoint in (d). (f) Percent fat mass change from baseline in mice treated with PT2399 or vehicle, measured by MRS body scan. (g, h) Scatter plots of inguinal (iWAT) and epididymal (eWAT) white adipose tissue mass versus pre-treatment body weight in non-tumor-bearing (NTB) controls and OSRC-2 xenograft-bearing mice treated with vehicle or PT2399. Each dot represents an individual mouse. Group-specific linear regression lines are shown. Analysis of covariance (ANCOVA) adjusting for pre-treatment body weight revealed significantly higher iWAT and eWAT mass in PT2399-treated mice compared to vehicle-treated mice (iWAT: p = 0.020, 95% CI: 0.02–0.15 g; eWAT: p = 0.008, 95% CI: 0.03–0.13 g). (i-k) (i) Immunofluorescent (IF) stain with DAPI and Ucp1 for iWAT, (j) hematoxylin and eosin (H&E) staining of iWAT and eWAT sections from NT, vehicle-treated, and PT2399-treated groups. (k) Gross anatomical images and isolated iWAT depots from NT, vehicle-treated, and PT2399-treated mice. For all panels, data are presented as mean ± SEM. Statistical significance was assessed using unpaired two-tailed t-tests. P values are indicated as follows: *P* ≤ *0.05* (\**), P ≤ 0.005 (\*\**), *P ≤ 0.005* (***), and *P* ≤ *0.0005* (****). (a) Created in https://www.BioRender.com.

To study this phenomenon further, we did MRS body scans of OSRC-2 tumor-bearing mice before and 17 days after treatment with PT2399 or vehicle. Once again, the vehicle-treated mice progressively lost weight, while PT2399-treated mice largely maintained their body weights despite comparable tumor burdens at necropsy (Extended Data Fig. 1e,f). Immunoblot assays further confirmed the PD effects of PT2399 (Extended Data Fig. 1g). Daily food intake did not differ between the two groups and was not diminished compared to non-tumor bearing mice (Extended Data Fig. 1h,i). Body scans indicated that the weight loss in vehicle-treated mice was primarily due to reductions in fat mass, with lean mass largely being preserved (Fig. 1f and Extended Data Fig. 1j). To more accurately assess adipose and skeletal muscle tissue wasting while addressing known limitations of normalizing tissue weight to body weight, we plotted fat mass against pre-treatment body weight and performed analysis of covariance (ANCOVA). This analysis showed that PT2399 treatment preserved inguinal and epididymal white adipose tissue (iWAT and eWAT) compared to vehicle-treated controls (Fig. 1g,h). Similarly, brown adipose tissue (BAT) and gastrocnemius muscle mass were also preserved in PT2399-treated mice (Extended Data Fig. 1k,l), while the assumption for ANCOVA was not met for quadriceps muscle (Extended Data Fig. 1m). The decreased WAT mass in vehicle-treated OSCR2 tumor-bearing mice was associated with the upregulation of the thermogenic protein Ucp1 (Fig. 1i and Extended Data Fig. 1n), which causes energy dissipation as heat (thermogenesis) rather than through ATP generation ^28^. Co-immunofluorescence of adipose tissue revealed that Ucp1, which marks white adipocytes that are converting to brown-like adipocytes with thermogenic properties (i.e. “browning”), was reduced in the adipose of PT2399-treated mice (Fig. 1i and Extended Data Fig. 1n). In addition to Ucp1 induction, white adipocytes in vehicle-treated OSRC-2 tumor-bearing mice exhibited the hallmark switch from unilocular lipid-storing white adipocytes to multilocular thermogenic brown/beige adipocytes, a process that was blocked upon PT2399 treatment (Fig. 1j,k and Extended Data Fig. 1o). Collectively, these results suggest that cachexia in this model is largely due to increased calorie utilization rather than decreased caloric intake and that PT2399 treatment blocks the induction of thermogenic gene programs in adipocytes.

To ask how, mechanistically, OSRC-2 cells induce cachexia, we introduced a promiscuous biotin ligase (BirA) fused to an endoplasmic reticulum targeting signal (ER) into OSRC-2 cells and, as a control, 786-O *VHL-/-* ccRCC cells. 786-O cells do not induce profound cachexia in mouse xenograft assays. The Bir-ER fusion has been used by others to biotinylate secreted proteins that can then be captured using streptavidin agarose (SA) and identified by mass spectrometry (MS) ^29^. In pilot experiments, we confirmed that cells expressing BirA-ER and treated with biotin secreted biotinylated proteins. To do so, proteins recovered from conditioned media using SA were eluted by boiling in SDS-containing sample buffer, resolved by SDS-PAGE, and transferred to nitrocellulose filters that were probed with streptavidin-horseradish peroxidase (Fig. 2a,b). Streptavidin blots of conditioned media showed that cells expressing BirA-ER biotinylated a broad range of secreted proteins compared with conditioned media from cells lacking BirA-ER. The background signal in the BirA-ER OSRC-2 cells in the absence of supplemental biotin likely reflects that these cells, in contrast to the BirA-ER 786-O cells, were grown in RPMI media that contains low levels of biotin. Immunoblotting confirmed that known secreted proteins IGBP3 and MMP2 were biotinylated in a BirA-ER dependent manner in both cell lines.

**Figure 2.**
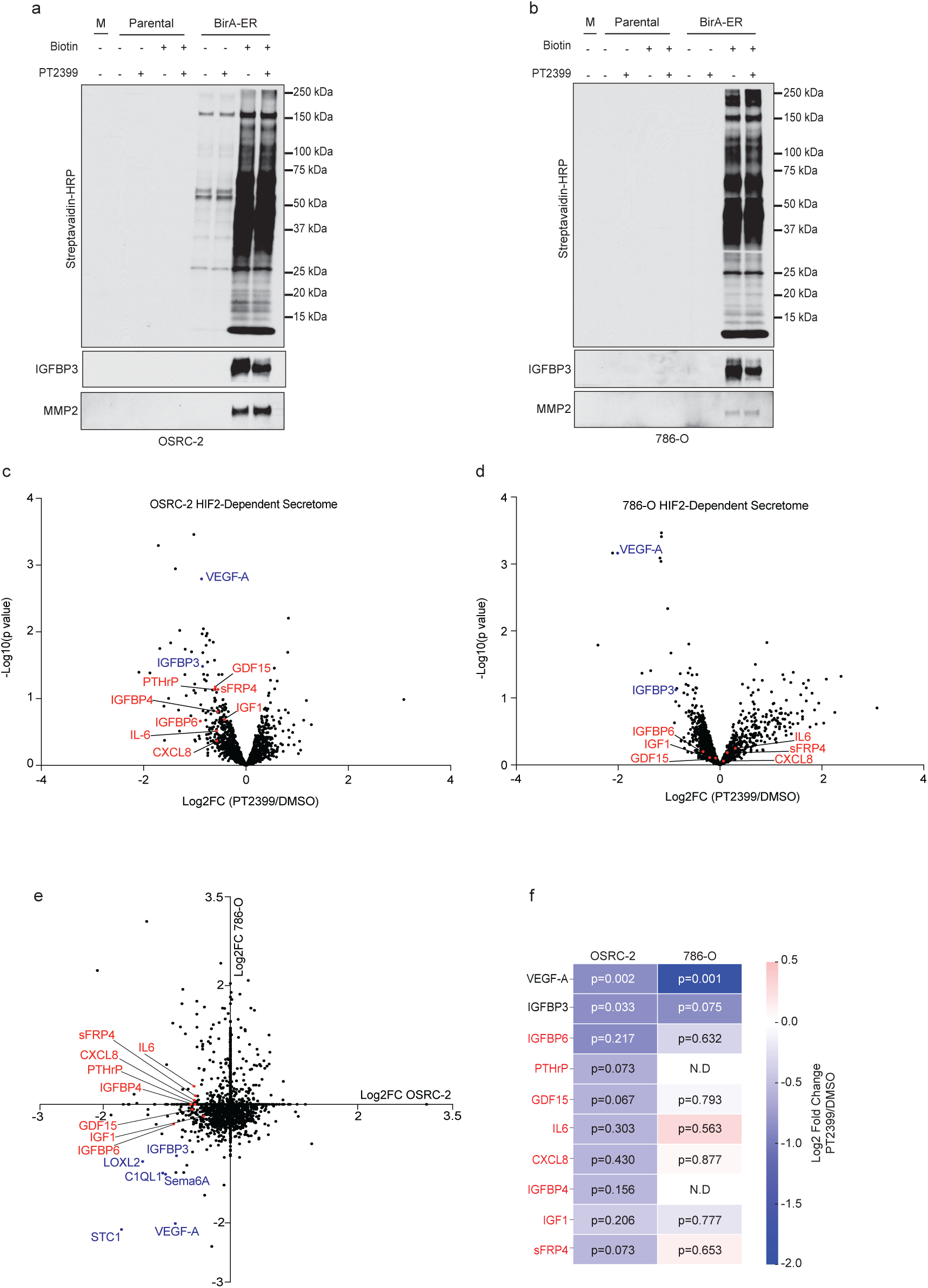
Identification of HIF2-responsive proteins secreted by OSRC-2 cells. (a and b) Parental and BirA-ER OSRC-2 cells (a) or parental and BirA-ER 786-O cells (b) treated, where indicated, with biotin and/or 2 mM PT2399 for 48 hours. Secreted proteins from conditioned media (or unconditioned media, “M”) were captured with streptavidin agarose, resolved by SDS-PAGE, transferred to nitrocellulose, and detected with streptavidin-HRP or by immunoblotting with anti-IGBP3 or anti-MMP2 antibodies. (c and d) Volcano plots of -log10(p-value) versus log2 fold change (Log2FC) for proteins secreted by OSRC-2 cells (c) or 786-O cells (d) treated with PT2399 compared to DMSO (n = 2). The known HIF2-regulated proteins VEGF-A and IGFBP3 are highlighted in blue and OSRC-2-specific downregulated proteins in red. (e) Scatter plot comparing Log2FC values for secreted proteins regulated by PT2399 in OSRC-2 vs 786-O cells, showing overlapping known HIF2-regulated proteins in blue and OSRC-2-specific downregulated proteins in red. (f) Heatmap displaying the log2 fold change for proteins secreted by OSRC-2 and 786-O cells treated with PT2399 vs DMSO control. P-values indicate statistical significance for each protein. ND = non-detected proteins. Proteins in red appeared to be more sensitive to PT2399 in OSRC-2 cells than in 786-O cells.

To identify biotinylated proteins secreted by OSRC-2 and 786-O cells, we treated these cells (parental), as well as BirA-ER versions of these cells, with PT2399 or DMSO for 48 hrs, captured secreted proteins with SA, and identified secreted proteins by LC-MS/MS using TMT (tandem mass tagging)-based quantification. We identified multiple PT2399-responsive secreted growth factors, including many that were shared across the two cell lines and some, such as VEGF-A, that are usually present in low (pM) abundance (Fig. 2c,d).

We focused on a short list of secreted proteins that were repressed by PT2399 in OSRC-2 cells but were not repressed, or far less repressed, by PT2399 in 786-O cells (proteins in red in Fig. 2e,f). Among these were PTHrP, GDF15, and IL-6, all of which have been linked to cachexia in various settings ^18,19,30,31^. We confirmed that *PTHLH* and its protein product, PTHrP, are repressed by PT2399 in OSRC-2 cells and highly expressed in OSRC-2 cells relative to 786-O cells (Extended Data Fig. 2a-d). In contrast, *GDF15* mRNA and secreted protein levels were not repressed by PT2399 in OSRC-2 cells (Extended Data Fig. 2a,e). Moreover, GDF15 levels in OSRC-2 cell conditioned media were comparable to the levels observed for 786-O cell conditioned media and normalized *GDF15* mRNA levels were higher in 786-O cells than in OSRC-2 cells (Extended Data Fig. 2d,e). *IL-6* mRNA levels were similarly not repressed by PT2399 (Extended Data Fig. 2a). We therefore focused on PTHrP.

*PTHLH* has been reported to be HIF-responsive in various settings, including ccRCC, and to have a cis-acting HIF binding site ^24,25,32,33^. We confirmed the presence of the latter in ChIP-Seq assays done with OSRC-2 cells wherein a FLAG-HA epitope tag was or was not appended to the endogenous HIF2α open reading frame using CRISPR homology-directed repair (HDR) ^27^ (Fig. 3a). We confirmed that *PTHLH* was amongst the most HIF2-responsive genes in OSRC-2 cells in steady-state RNA-Seq experiments done with OSRC-2 cells treated with PT2399 (Fig. 3b and Extended Data Fig. 3a,b) or after CRISPR-KO of *EPAS*1 (Fig. 3c), which encodes HIF2α. We validated these RNA-Seq findings by quantitative RT-PCR (Extended Data Fig. 3c). In Pro-Seq experiments *PTHLH* transcription was dramatically reduced, as determined by RNA Pol II recruitment, within 2 hours of PT2399 treatment (Fig. 3d,e and Extended Data Fig. 3d). Polysome-seq analysis of OSRC-2 cells treated with PT2399 for 72 hours revealed significant translational downregulation of *PTHLH* (Fig. 3f). Therefore, *PTHLH* appears to be a direct HIF2 target in OSRC-2 cells.

**Figure 3.**
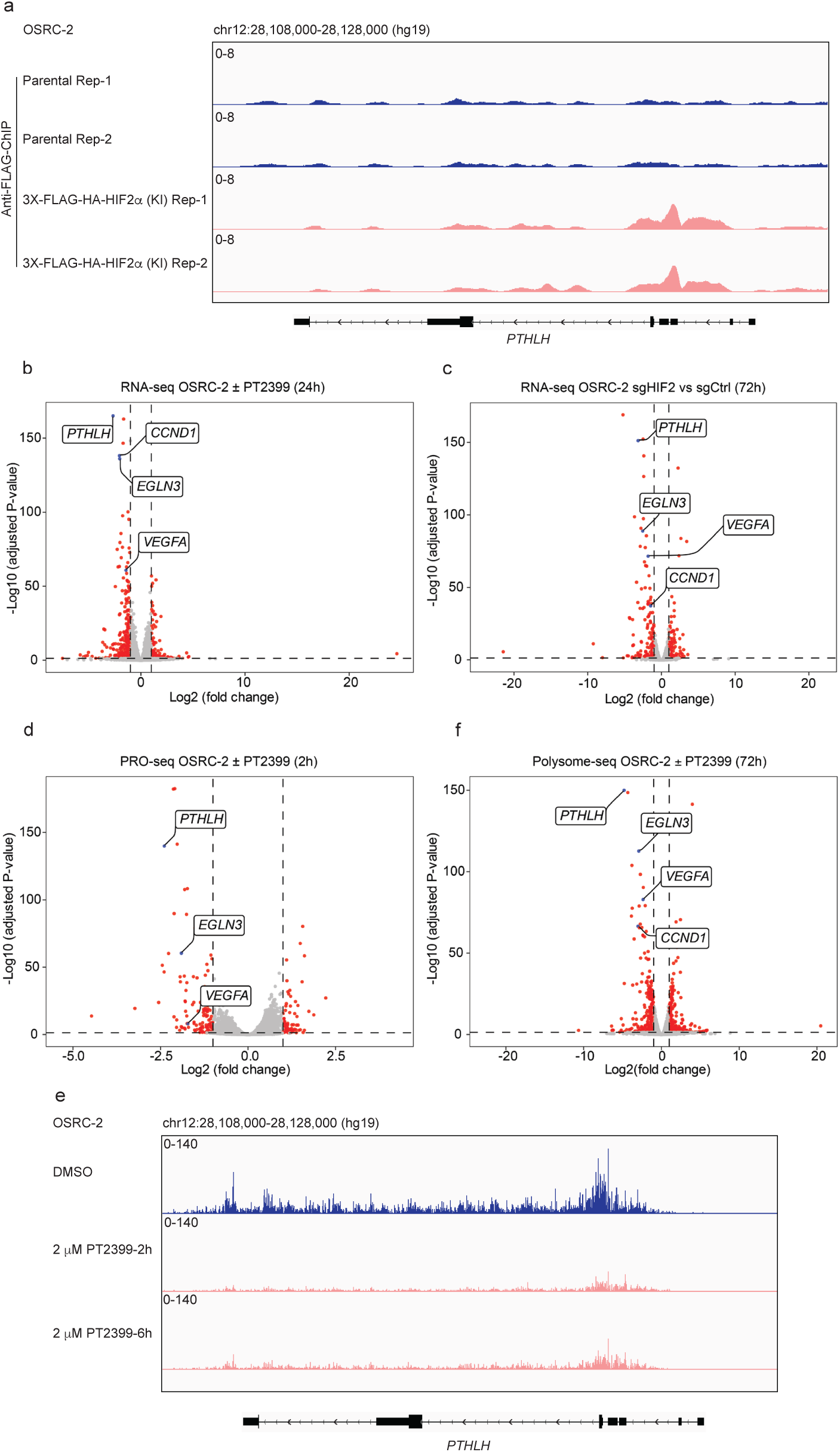
*PTHLH* is a direct HIF2 target gene in OSRC-2 cells. (a) Anti-Flag ChIP-seq tracks (2 biological replicates) at the *PTHLH* locus in OSRC-2 cells with parental and 3X-FLAG-HA-HIF2α knock-in (KI) cells. The y-axis represents ChIP-seq read intensity, and the x-axis corresponds to the chromosomal region chr12:28,108,000– 28,128,000 based on the hg19 human genome assembly. (b and c) Volcano plots of RNA-seq data comparing OSRC-2 cells treated with 2 mM PT2399 or DMSO for 24 hours (b) or infected to express sgHIF2α or sgCtrl (c). *PTHLH* and selected HIF2 target genes are highlighted in blue. (d) Volcano plot of PRO-seq data from OSRC-2 cells treated with 2 mM PT2399 or DMSO for 2 hours. (e) PRO-seq tracks at the *PTHLH* locus in OSRC-2 cells treated with DMSO or 2 mM PT2399 for 2 and 6 hours. The y-axis represents transcriptional read intensity, and the x-axis corresponds to the chromosomal region chr12:28,108,000–28,128,000 based on the hg19 human genome assembly. (f) Volcano plot of polysome-seq data from OSRC-2 cells treated with 2 mM PT2399 or DMSO for 72 hours. Data are presented as Log2 fold change versus -Log10 adjusted P value. For volcano plots, statistical thresholds for significance were set at P ≤ 0.05, and n = 3. The adjusted p-value for *PTHLH* was zero but was set slightly above the smallest non-zero adjusted p-value of the other genes for graphical purposes.

Hypercalcemia driven by PTHrP has been described in other cancers, including squamous cell cancers of the head, neck, and lungs ^34^. We observed that *PTHLH* expression in the head and neck cancer cell line SCC-9 was induced by hypoxia (Extended Data Fig. 3e,f). In this setting, HIF1α and HIF2 cooperatively regulated *PTHLH* because *PTHLH* expression in SCC-9 was blocked by CRISPR-based inactivation of ARNT or by treatment with PT2399 if they were engineered to also lack HIF1α (Extended Data Fig. 3g,h). Qualitatively similar results were seen with a second head and neck cancer line, SCC-4 (Extended Data Fig. 3i-l). Therefore, HIF likely regulates *PTHLH* in cancers beyond kidney cancer.

We next performed loss of function and gain of function *PTHLH* studies to ask if PTHrP was necessary or sufficient, respectively, for the rapid induction of cachexia by OSRC-2 cells. For the former, we performed orthotopic tumor assays with polyclonal OSRC-2 cells stably infected with a lentivirus expressing 1) Cas9, 2) firefly luciferase, and 3) either a *PTHLH* sgRNA or a control (AAVS1) sgRNA. Tumor burden was monitored by serial bioluminescent imaging (BLI). CRISPR-mediated knockout of *PTHLH* in OSRC-2 cells was verified by amplicon sequencing and dramatically decreased PTHrP secretion *in vitro* (Fig. 4a and Extended Data Fig. 4a). *PTHLH* mRNA levels, but not other HIF2-responsive mRNA levels, were modestly reduced in the CRISPR *PTHLH* knockout (KO) cells, presumably due to mRNA destabilization resulting from indel mutations (Extended Data Fig. 4b-d).

**Figure 4.**
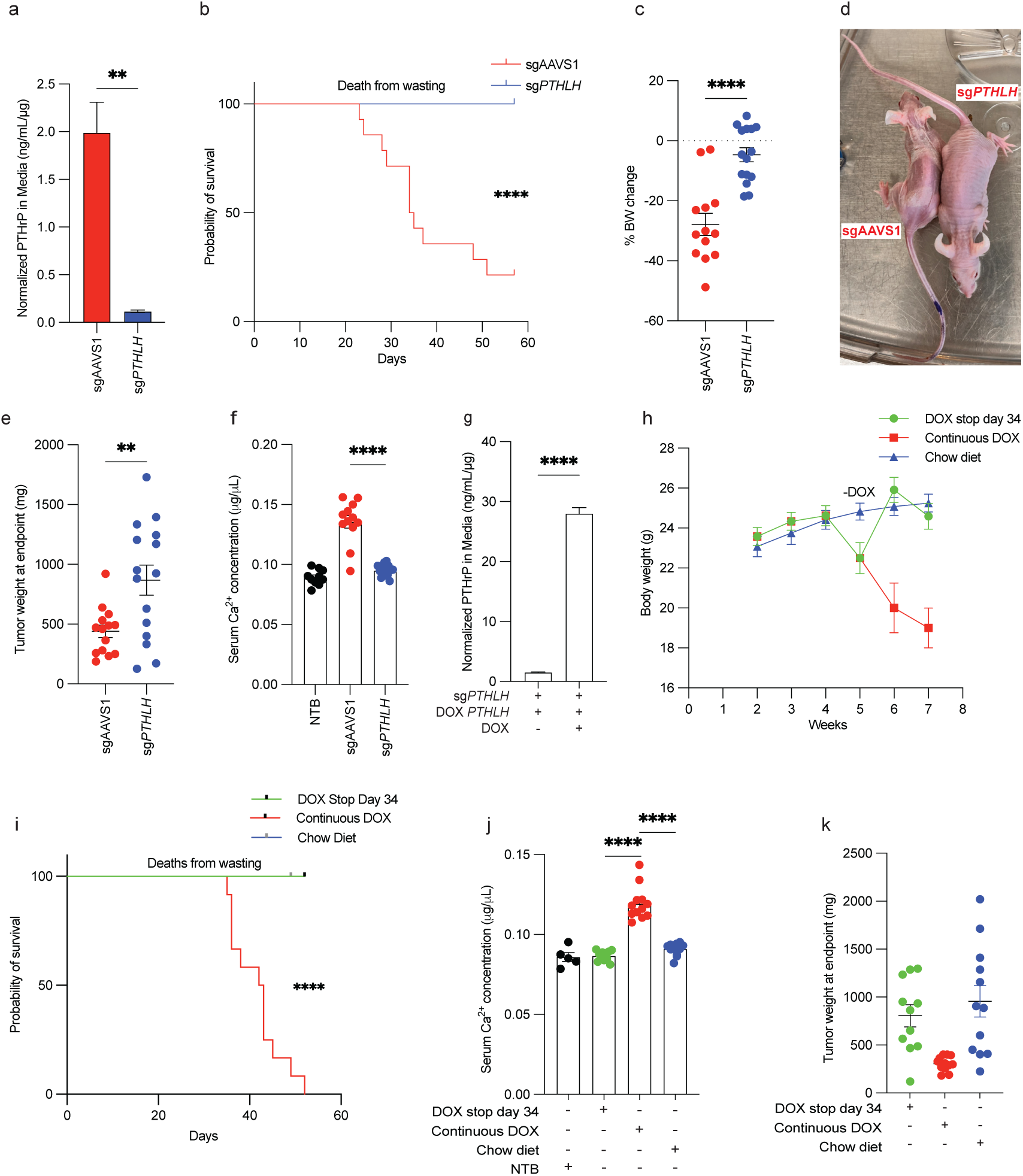
*PTHLH* is necessary for the induction of cachexia by OSRC-2 tumors. (a) PTHrP levels in conditioned media from OSRC-2 cells infected to express sg*PTHLH* or sgAAVS1, normalized to total cellular protein (n = 3). (b) Kaplan-Meier survival curve for mice with OSRC-2 tumors infected to express sg*PTHLH* (blue, n = 15) or sgAAVS1 (red, n = 14). (c-f) Body weight changes (c), representative mouse images (d), tumor weights (e), and serum calcium levels (f) at study endpoint of mice from (b). NTB = non-tumor bearing. (g) PTHrP levels in conditioned media from OSRC-2 sg*PTHLH* cells infected to express a DOX-inducible, sgRNA-resistant, *PTHLH* cDNA and grown in the presence or absence of doxycycline (DOX), normalized to total cellular protein (n = 3). (h-i) Body weights (h) and Kaplan-Meier survival curves (i) of mice bearing tumors formed by OSRC-2 cells from (g). The mice were continuously fed DOX, never fed DOX (Chow diet), or had DOX stopped 34 days after cell implantation. (j and k) Serum calcium (j) and tumor weights (k) at the endpoint in (i). Statistical analysis for Kaplan-Meier survival curve was performed using the log-rank test. For other panels, data are presented as mean ± SEM. Statistical significance was assessed using unpaired two-tailed t-tests. P values are indicated as follows: *P* ≤ *0.05* (\**), P ≤ 0.005 (\*\**), *P ≤ 0.005* (***), and *P* ≤ *0.0005* (****).

Mice implanted with OSCR-2 cells edited with the AAVS1 sgRNA developed cachexia and were sacrificed when moribund. In contrast, the mice implanted with the OSRC-2 cells that had undergone CRISPR-KO of *PTHLH* did not lose weight and were not sacrificed until the last control mice needed to be euthanized (day 57) (Fig. 4b-d). The sg*PTHLH* tumors were significantly larger than the sgAAVS tumors (Fig. 4e), arguing that the absence of cachexia with the former was not because they were insufficiently large. Note that the sgAAVS tumors had to be excised earlier due to cachexia and hence this experiment does not indicate that *PTHLH* is a ccRCC suppressor. Serum PTHrP was below the sensitivity limits of the commercial ELISA kits we tested. However, PTHrP has also been implicated in humoral hypercalcemia ^35^. Serum calcium levels were increased mice with sgAAVS1 tumors compared to the mice with sg*PTHLH* tumors and to non-tumor bearing mice (NTB), suggesting that PTHrP was elevated in mice with sgAAVS1 tumors (Fig. 4f).

To ask if the *PTHLH* KO phenotype was on-target, we superinfected the *PTHLH* KO OSRC-2 cells with a lentivirus expressing a doxycycline (DOX), sgRNA-resistant, *PTHLH* cDNA that encoded PTHrP with a C-terminal V5 epitope tag (Fig. 4g and Extended Data Fig. 4e). We confirmed that inducible *PTHLH* expression did not affect other canonical HIF2-responsive mRNAs (Extended Data Fig. 4e,f). Mice orthotopically implanted with these cells and continuously fed DOX-containing chow, but not normal chow, developed hypercalcemia and cachexia about 4 weeks later (Fig. 4h-j and Extended Data Fig. 4g). Notably, removing DOX in the former mice, thus silencing *PTHLH*, normalized serum calcium, reversed the cachexia and prolonged survival (Fig. 4h-j and Extended Data Fig. 4g). These effects were again not explained by differences in tumor burden measured at the study endpoint (Fig. 4k). To determine whether hypercalcemia was the underlying cause for the cachectic phenotype, we treated OSRC-2 tumor-bearing mice with the calcium-lowering bisphosphonate zoledronic acid ^36^ (ZOL; 120 µg/kg intraperitoneally every other day), with the HIF2 inhibitor PT2399 (45 mg/kg daily oral gavage), or with vehicle (Extended Data Fig. 4h-k). As before, vehicle-treated mice developed progressive weight loss and hypercalcemia, whereas PT2399 reversed both phenotypes independently of tumor weight. ZOL rapidly normalized serum calcium within 3-4 days of treatment initiation (Extended Data Fig. 4l-n), but did not reverse the cachexia. As a result, ZOL-treated mice ultimately succumbed to severe cachexia despite correction of hypercalcemia. These findings indicate that hypercalcemia is not necessary for the cachexia we observe in this model.

To address sufficiency, we infected 786-O with a lentivirus expressing a DOX-inducible *PTHLH* cDNA that also encodes a 3’ V5 epitope tag (DOX-*PTHLH*) or, as a control, the empty vector (EV). Treatment of the DOX-*PTHLH* cells increased PTHrP secretion *in vitro*, as expected, again without affecting other HIF2 target genes (Fig. 5a and Extended Data Fig. 5a-c). Next the DOX-*PTHLH* cells and EV cells were orthotopically implanted in nude mice fed normal chow. Tumor formation was monitored by serial BLI. Six weeks after implantation, the tumor bearing mice were randomized to DOX-containing chow or normal chow. Mice with DOX-*PTHLH* tumors, but not EV tumors, rapidly developed cachexia and hypercalcemia, requiring euthanasia within several weeks, if fed the DOX-containing chow (Fig. 5b-e and Extended Data Fig. 5d). The tumors removed from these euthanized mice were not larger than tumors removed from the EV and no DOX control groups at the time the last DOX-*PTHLH* tumor bearing mice needed to be sacrificed (day 56) (Fig. 5f).

**Figure 5.**
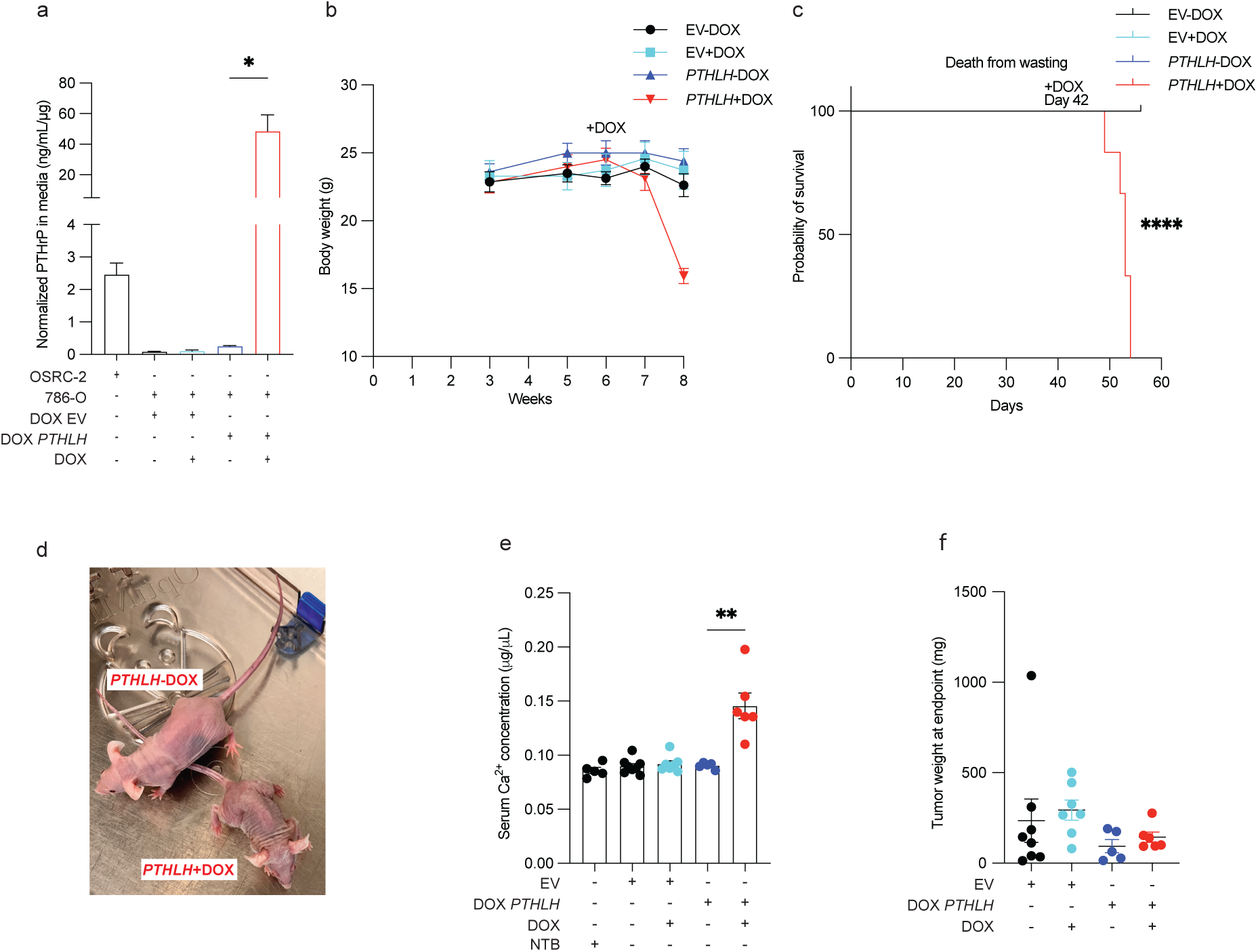
*PTHLH* is sufficient for the induction of cachexia by 786-O xenografts. (a) PTHrP levels in conditioned media from 786-O cells infected with doxycycline (DOX)-inducible *PTHLH* lentivirus or empty vector (EV) and grown in the presence or absence of DOX, normalized to total cellular protein (n = 3). (b and c) Body weights (b) and Kaplan-Meier survival curve (c) for mice with xenografts formed by 786-O cells from (a) and maintained on chow with or without DOX. Statistical analysis was performed using the log-rank test (****P ≤ 0.0001). (d-f) Representative mouse images (d), serum calcium levels (e), and tumor weights (f) at study endpoint of mice from (b and c). Statistical significance was assessed using unpaired two-tailed t-tests. P values are indicated as follows: *P* ≤ *0.05* (\**), P ≤ 0.005 (\*\**), *P ≤* 0.005 (***), and *P* ≤ *0.0005* (****).

To ask about the generalizability of our preclinical findings, we surveyed *PTHLH* expression in 7 additional ccRCC cell lines. RFX393 cells had the second highest *PTHLH* mRNA levels (after OSRC-2 cells) and secreted measurable amounts of PTHrP (Extended Data Fig. 6a,b), both of which were suppressed by PT2399 (Extended Data Fig. 6c,d). As was true for OSRC-2 cells, RXF393 cells induced cachexia and hypercalcemia in subcutaneous xenograft assays and these two phenotypes were reversed by PT2399 (Extended Data Fig. 6e-h) and prevented by CRISPR KO of *PTHLH* without commensurate changes in tumor growth (Extended Data Fig. 6i-n). Therefore, upregulation of PTHrP by HIF2 causes cachexia and hypercalcemia in both the OSRC-2 and RXF393 models.

*PTHLH* resides on chromosome 12p in a region that is amplified in different histologic types of kidney cancer, including ccRCC ^37^, as well as in other cancers, including pancreatic ductal adenocarcinoma^38^. Notably, an analysis of available ccRCC cell lines from DepMap ^39^ revealed that the copy number of this region is increased in OSRC-2 cells compared to 786-O cells and that *PTHLH* expression positively correlates with *PTHLH* copy number across ccRCC cell lines (Extended Data Fig. 7a). Using TCGA data, we confirmed that *PTHLH* is frequently highly expressed in human ccRCC, especially in those that have gained or amplified chromosome 12p (Extended Data Fig. 7b-e). We performed a comprehensive pan-cancer transcriptomic analysis and found that *PTHLH* expression strongly correlates with a HIF transcriptional signature across multiple tumor types, with particularly strong associations in head and neck squamous cell carcinoma (HNSC) and lung squamous cell carcinoma (LUSC) (Extended Data Fig. 7f-m). Moreover, TCGA analysis revealed that HNSC exhibits the highest overall *PTHLH* mRNA expression levels among the tumor types we examined (Extended Data Fig. 7b).

To begin to address the clinical utility of our findings, we studied ccRCC patients treated with the allosteric HIF2 inhibitor belzutifan for whom appropriate plasma samples and clinical information were available. Comparable patients treated with an immune checkpoint inhibitor (ICI), or a VEGF receptor tyrosine kinase inhibitor (TKI) served as controls, with the caveat that none of the belzutifan patients received belzutifan as their first line of therapy, in contrast to most of the ICI and TKI patients, where the ICIs and TKIs were frequently used as front line therapy (Extended Data Table 1). Plasma PTHrP was measured in a Mayo Clinic CLIA Laboratory ^40^.

We identified 26 patients with plasma samples before and after belzutifan, 36 patients with plasma samples before and after ICI therapy and 38 patients with samples before and after TKI therapy. We confirmed that plasma PTHrP was elevated in the ccRCC patients compared to 12 age-matched healthy volunteers (Fig. 6a). Pre-treatment PTHrP levels positively correlated with the pre-treatment corrected calcium levels (Fig. 6b). In this cohort, patients with baseline PTHrP levels above the median had significantly lower baseline skeletal muscle index compared to those with lower PTHrP levels (Fig. 6c). Belzutifan, but not the ICI or VEGF TKI, suppressed plasma PTHrP (Fig. 6d and Extended Data Fig. 8a,b) and corrected calcium after 1 month of treatment (Fig. 6e). Notably, the belzutifan treated patients gained weight during this period, while weight was unchanged in the ICI group and fell in the VEGF TKI group (Fig. 6f and Extended Data Fig. 8c). The same pattern emerged when this analysis was restricted to patients with pre-treatment BMIs < 30 (Extended Data Fig. 8d,e). These effects of belzutifan did not appear to be a secondary consequences of tumor shrinkage and loss of viable tumor cells because 1) they occurred rapidly, 2) were observed in both belzutifan responders and non-responders, and 3) the clinical response rate to belzutifan, if anything, trended lower than the response rates to ICI and VEGF TKI in our cohorts (Extended Data Fig. 8f and Extended Data Table 1).

**Figure 6.**
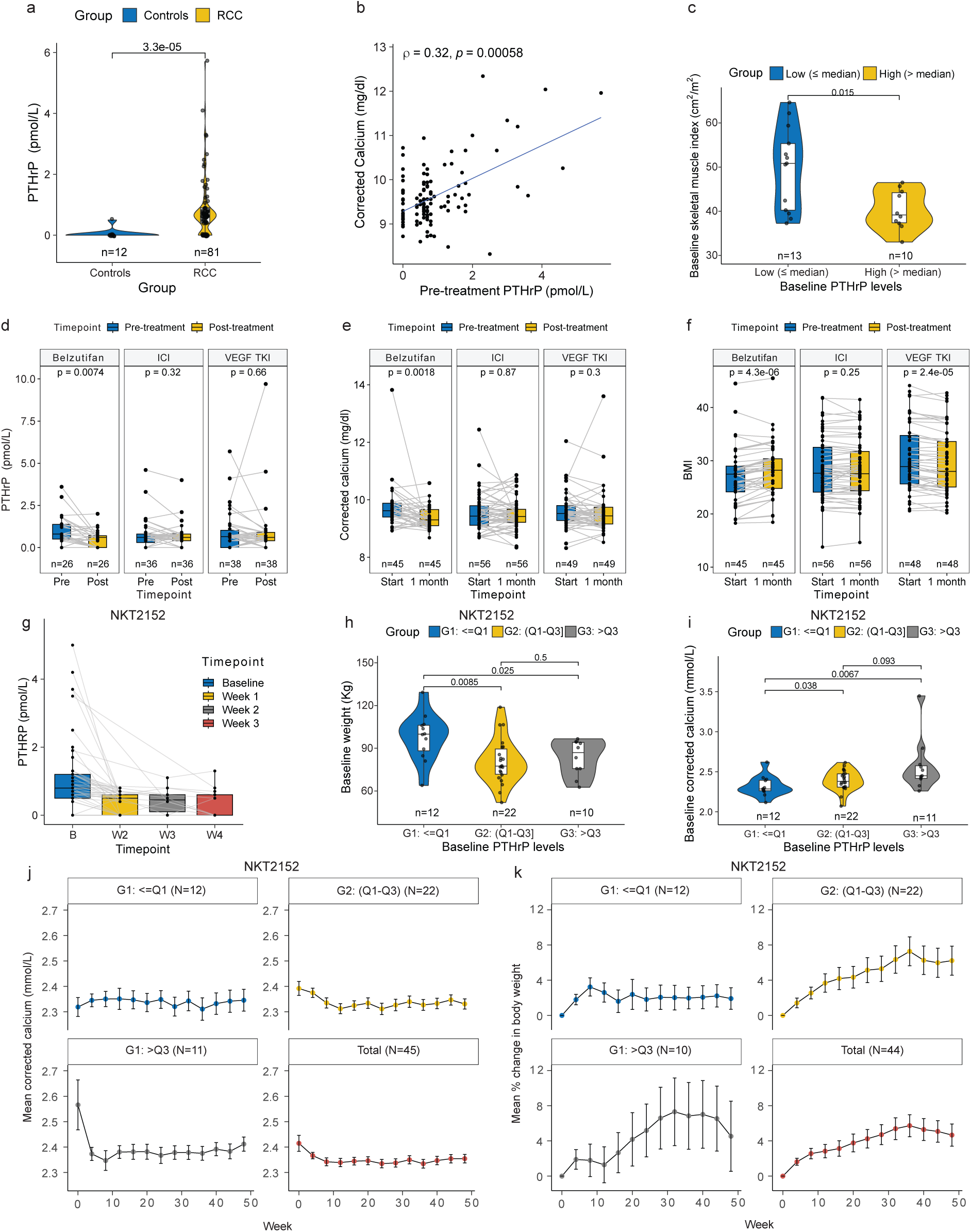
Treatment of kidney cancer hypercalcemia and cachexia with HIF2 inhibitors. (a) Plasma PTHrP levels in untreated healthy controls (blue, n = 12) and RCC patients (yellow, n = 81). (b) Correlation between pre-treatment plasma PTHrP levels and pre-treatment albumin-corrected calcium levels (n = 112). (c) Skeletal muscle index (SMI; cm²/m²) at baseline in patients stratified by circulating PTHrP levels (n = 23). Patients were divided into low (≤ median, 0.8 pmol/mL; n = 13, blue) and high (> median; n = 10, yellow) PTHrP groups. (d-f) Plasma PTHrP levels (d), corrected serum calcium levels (e), and body mass index (BMI) (f) in patients treated with belzutifan (n = 26 for b, n = 45 for c, n = 43 for d), immune checkpoint inhibitors (ICIs; n = 36 for b, n = 56 for c, n = 56 for d), or VEGF tyrosine kinase inhibitors (TKIs; n = 38 for b, n = 49 for c, n = 48 for d), pre-treatment and post-treatment. (g) Plasma PTHrP levels over time (baseline, Week 1, Week 2, and Week 3) in RCC patients treated with the HIF2 inhibitor NKT2152 (n = 45). (h) Baseline body weight of RCC patients stratified by plasma PTHrP levels: low (<Q1, n = 12), mid (Q1-Q3, n = 22), and high (>Q3, n = 10). (i) Baseline albumin-corrected calcium levels stratified by baseline plasma PTHrP levels: low (<Q1, n = 12), mid (Q1-Q3, n = 22), and high (>Q3, n = 11). (j and k) Mean corrected serum calcium (j) and mean percent change body weight (k) over 50 weeks in RCC patients from (h) treated with NKT2152, stratified by plasma PTHrP levels. Windowing algorithm and Last Observation Carried Forward (LOCF) were applied for plotting (j) and (k). Data in (j and k) are shown for patients with low (<Q1, n = 12), mid (Q1-Q3, n = 22), and high (>Q3, n = 11 for j, n = 10 for (k) due to one subject missing baseline body weight information) baseline PTHrP levels. Data are presented as mean ± SEM. Statistical significance was assessed for (a), (c), (h), and (i) by using unpaired Wilcoxon rank sum test, for (d-f) using the paired Wilcoxon signed-rank test, and for (b) using Spearman’s rank correlation coefficient.

To further probe the robustness of these findings, we obtained plasma samples and clinical data for 45 ccRCC patients treated with the allosteric HIF2 inhibitor NKT2152, which is currently under clinical development ^41^. NKT2152 caused a rapid and sustained suppression of PTHrP (Fig. 6g and Extended Data Fig. 8g). We then subdivided the patients based on their pre-treatment PTHrP values as follows: Group 1, the 12 patients with the lowest PTHrP levels, Group 2, the 22 subjects with baseline PTHrP levels in the 25th to 75th percentile of PTHrP levels, and Group 3, the 11 subjects with the highest PTHrP levels. A higher proportion of Group 3 patients had Stage IV disease, presumably reflecting the anticipated correlation between PTHrP levels and total tumor burden (Extended Data Table 2). It is also known that high *PTHLH* levels correlate with worse outcomes in ccRCC ^42^. As expected from our preclinical findings, Group 3 had the lowest body weights and highest corrected serum calcium levels of the three groups at baseline (Fig. 6h,i) and appeared to derive the greatest benefit from NKT2152 treatment with respect to these two measures (Fig. 6j, k). Note that one group 3 patient was omitted from Fig. 6h,k because they did not have a baseline body weight value. In contrast, the clinical response rates did not appear to be higher in Group 3 compared to the other two groups (Extended Data Table 2). As was true for belzutifan, weight gain in patients treated with NKT-2152 was observed even in patients with stable or progressive disease, indicating that cachexia reversal can occur independently of tumor shrinkage (Extended Data Fig. 8h). Sex-disaggregated analyses for belzutifan-treated patients (Extended Data Table 1) and NKT2152-treated patients (Extended Data Table 2), based on sex assigned at birth, are provided in the source data files and showed that key trends were similar in males and females. Therefore, two different allosteric HIF2 inhibitors appear to mitigate two of the common paraneoplastic syndromes associated with ccRCC.

## Discussion

Paraneoplastic syndromes are common in ccRCC patients and often complicate their management ^12–14^. In this study, we strengthened the earlier conclusion that *PTHLH* is a direct HIF2 target gene ^24,25,32,33^, although our findings do not exclude the possibility that pVHL loss also stabilizes the *PTHLH* mRNA, as suggested by others previously ^43^. We provide further evidence that the *PTHLH* gene product, PTHrP, causes the hypercalcemia that frequently complicates ccRCC ^42,44–46^. More importantly, we provide preclinical and clinical evidence that PTHrP also causes cachexia that frequently complicates ccRCC and that both hypercalcemia and cachexia in ccRCC can be managed with pharmacological HIF2 inhibitors. Notably, not all ccRCC patients develop hypercalcemia and cachexia ^14^. The development of these two ccRCC complications is likely to be influenced by multiple factors, including the degree of HIF dysregulation, *PTHLH* copy number variation, and the epigenetic accessibility of cis-regulatory HIF-binding elements (e.g. whether they are unmethylated). Although hypercalcemia can, itself, cause nausea, vomiting, and decreased appetite, the cachexia in our mouse models was not associated with decreased caloric intake. It was instead associated with adipocyte changes previously shown to be mediated by the PTHrP receptor ^18,19^.

PTHrP facilitates cachectic wasting through the engagement with its cognate receptor, Parathyroid Hormone 1 Receptor (PTH1R), on adipocytes ^18,19,47,48^. Consistent with prior results, we found that PTHrP leads to adipose tissue wasting and causes a switch from an energy-storing phenotype to an energy-consuming phenotype in adipocytes, as marked by Ucp1 induction. Notably, the role of Ucp1-mediated browning in driving cachectic wasting remains controversial. To test the functional role of Ucp1 in wasting, whole body *Ucp1* deletion animals (Ucp1^-/-^) have been used, which demonstrated that Ucp1 loss blocks cachectic wasting in some contexts^49^, but not others^50^. Thus, Ucp1-independent mechanisms, such as increased lipolysis or decreased lipogenesis, have been suggested to also contribute to cachexia. While we noted increased Ucp1 expression in cachectic adipose, we did not test its functional role in driving wasting, and it remains to be explored if lipolytic and lipogenic pathways also contribute to PTHrP-mediated wasting in this context. Tumor-derived PTHrP has recently been shown to downregulate *de novo* lipogenesis (DNL) in adipose tissue from mouse models of pancreatic cancer-associated cachexia^48^. Furthermore, patients with high circulating PTHrP demonstrate increased weight loss and whole body fat oxidation independent of peripheral lipolysis^23^. The interplay between browning, lipogenesis, and lipolysis during cachexia is complex, and the role of PTHrP in regulating these processes in RCC-mediated wasting warrants further study. Notably, cachexia occurred rapidly in the OSRC-2 model of ccRCC cachexia, which necessitated early euthanasia of the tumor-bearing mice. At the early time points we were able to study in this model, adipose tissue wasting predominated, suggesting that this model reflects an early stage of cachexia. Nonetheless, ANCOVA adjusting for pre-treatment body weight revealed higher gastrocnemius mass in PT2399-treated mice compared to vehicle, whereas quadriceps showed a treatment-by-body weight interaction that precluded a definitive conclusion. Thus, skeletal muscle involvement is detectable but modest at these time points and is mitigated by HIF2 inhibition, while adipose tissue remains the principal early target. Supporting the clinical relevance of this observation, patients with baseline PTHrP levels above the median had significantly lower skeletal muscle index at baseline compared to those with lower PTHrP levels, consistent with a link between elevated PTHrP and reduced muscle mass^19^. As such, PTHrP-targeted interventions, either directly through anti-PTHrP therapies ^38,51,52^ or indirectly by targeting the HIF2-PTHrP axis, might serve to block the onset of cachexia.

*PTHLH* and PTHrP are downregulated acutely with HIF2 inhibitors and thus can be viewed as pharmacodynamic markers. Downregulation of HIF2 activity is presumably necessary, although not necessarily sufficient, for the antitumor effects of HIF2 inhibitors in ccRCC, as evidenced by the OSRC-2 model. Moving forward it will be important, as larger clinical datasets with appropriate biospecimens become available, to ask if elevated baseline PTHrP values and/or acute downregulation of PTHrP after treatment are positive predictive biomarkers for achieving disease control, including RECIST (response evaluation criteria in solid tumors) responses, with HIF2 inhibitors. Earlier work implicated PTHrP as a therapeutic target in ccRCC ^53^, although in our model, genetic ablation of *PTHLH* did not impede tumor growth. A caveat, however, is that these earlier studies used neutralizing antibodies and antagonist peptides that would affect both tumor-derived and host-derived PTHrP, in contrast to our studies ^38,53–56^.

In our preclinical models, reversal of cachexia with HIF2 inhibition occurred rapidly and could be dissociated from changes in tumor mass. We found that patients treated with HIF2 inhibitors tended to gain weight on therapy, in contrast to patients treated with standard of care agents such as ICIs or VEGF receptor TKIs. These findings are consistent with suppression of PTHrP reversing both clinically occult and clinically overt cachexia physiology. A caveat, however, is that our clinical data with respect to weight loss across treatment groups could be confounded by therapy-induced toxicities caused by tyrosine kinase inhibitors and ICIs, as well as by differences in patient characteristics across the different treatment groups, each of which was comprised of a relatively small number of patients for whom we had access to appropriate samples. Clearly it will be important to confirm our findings in larger studies. It will also be of interest to determine if concurrent treatment with belzutifan mitigates the risk of weight loss in patients treated with other ccRCC therapies, including VEGF TKIs.

The BirA-ER system, combined with quantitative LC-MS/MS, appears to be a robust engine for the discovery of proteins that are differentially secreted by cells under different conditions, such as, in our case, the presence or absence of a HIF2 inhibitor. We discovered multiple known HIF-responsive growth factors, including IGFBP3, PTHrP, and VEGF-A ^57^. It is likely that HIF2-dependent secreted proteins play roles in some of the other kidney cancer paraneoplastic syndromes for which the mechanism remains unknown. Although we focused on secreted proteins that were repressed by PT2399, secreted proteins that are induced after HIF2 blockade could also provide insights into HIF2 biology and lend themselves to serve as PD markers that would rise (“up assay”), rather than fall, in serum when HIF2 is successfully inhibited.

It will be important to determine if intratumoral hypoxia and HIF induce PTHrP in other cancers, such as pancreatic cancers ^23^, head and neck cancers ^58^, lung cancer ^18^ and colorectal cancer ^59^, where PTHrP has been implicated in cachexia and, if so, to determine which HIF paralog is responsible. In this regard, our pan-cancer analyses suggest that *PTHLH* is significantly correlated with HIF transcriptional activity in multiple cancer types, including HNSC and LUSC. Moreover, our HNSC cell line data suggest that both HIF1α and HIF2 drive *PTHLH* expression in this setting. Notably, the drug-binding pocket for HIF2α is much smaller than the corresponding pocket in HIF1α, which has thus far precluded the development of HIF1α inhibitors analogous to belzutifan and NKT2152 ^60^. Nonetheless, a number of drugs that directly or indirectly inhibit HIF1α have been described ^61^.

Our findings highlight an important role for PTHrP in ccRCC cachexia. In other settings, GDF15 contributes to cancer cachexia, and it is conceivable that both secreted factors are operative in some settings. An anti-GDF15 monoclonal antibody, ponsegromab, was recently shown to ameliorate cachexia caused by non-small cell lung cancer and pancreatic cancer ^62^, and CLIA-certified assays exist for measuring PTHrP and GDF15 in plasma. We envision the use of such assays in the future to tailor the use of PTHrP antagonists (direct or indirect via HIF) and GDF15 antagonists to treat cachexia.

## Methods

### Cell lines

OSRC-2 cells were obtained from the Riken Cell Bank and RXF393 cells were obtained from the National Cancer Institute (DCTD Tumor Repository). Both cell lines were maintained in RPMI 1640 (GIBCO; 11875093) supplemented with10% Fetal Bovine Serum (FBS) (GeminiBio; 100-106),100 U/mL Penicillin, and 100 μg/mL Streptomycin (P/S) (Gibco; 25200056). 786-O cells and 293T cells were obtained from the Kaelin Laboratory stocks and were maintained in DMEM (GIBCO; 11965092) supplemented with 10% FBS and P/S as above. SCC-9 and SCC-4 cells were obtained from American Type Culture Collection (ATCC) and were maintained in DMEM/F-12, GlutaMAX (GIBCO; 10565018) supplemented with 400 ng/mL hydrocortisone (Sigma-Aldrich; H0135-1MG) and 10% FBS and P/S as above. Lentiviral infected cells were maintained as polyclonal cells under drug selection. OSRC-2 and 786-O cells were selected with 800 μg/mL of G418, 10 μg/mL Blasticidin, or 2 μg/mL Puromycin. RXF393 cells were selected with 400 μg/mL of G418 or 2 μg/mL Puromycin. All cell lines were confirmed to be mycoplasma-free using a MycoAlert Mycoplasma Detection kit (Lonza; LT07-318). The OSRC-2, RXF393, 786-O, 293T, SCC-9 and SCC-4 cell lines used in this study were authenticated by short tandem repeat (STR) profiling. Authentication was performed by the Cell Line Authentication Service at the ATCC.

### Plasmids

The sgRNA-resistant PTHLH cDNA was created by site-directed mutagenesis using an Agilent QuikChange II XL Site-Directed Mutagenesis Kit (Agilent; 200522) according to the manufacturer’s instructions on a pENTR221 plasmid containing a wild-type *PTHLH* cDNA (Horizon Discovery; OHS5894-202497874). The primers used for site-directed mutagenesis and subcloning are listed below. The wild-type and mutant *PTHLH* cDNAs from the respective pENTR221 plasmids were then shuttled into pLIX_403-Puro vector (Addgene; 41395) or pLIX403-ccdB-Blast (Addgene; 158560) by using Gateway LR Clonase II Enzyme mix (Invitrogen; 11791100) according to the manufacturer’s instructions. The lentiCRISPR v2-Puro (Addgene; 52961), lentiCRISPR v2-Blast (Addgene; 83480), psPAX2 (Addgene; 12260), and pMD2.G (Addgene; 12259) were obtained from Addgene. The pLL3.7_EF1a_Fluc-neo vector was obtained from the Kaelin Laboratory stocks. The pLX304-BirA-G3*-ER construct was generated by recombining the BirA-G3*-ER ORF cDNA from a pDONR gateway entry clone (a gift from Norbert Perrimon’s Laboratory, Harvard Medical School) into the pLX304 vector (Addgene; 25890) using the Gateway cloning methodology as described above. The sgRNA expression vectors were digested with BsmBI (Thermo; ER0451) and ligated to annealed oligonucleotides encoding the desired sgRNAs using T4 ligase (NEB; M0202M) as described in ref ^63^. All *PTHLH* cDNA and sgRNA inserts were validated by DNA Sanger sequencing.

### Targeted mutagenesis primers

*PTHLH* sg*PTHLH* Resistant Forward 5′-CAGATGGTGAAGGAAGAACCGTCGCCGCAAGTCCTGAATGGACTTCCCCTTGTCA T-3′

*PTHLH* sg*PTHLH* Resistant Reverse 5′-ATGACAAGGGGAAGTCCATTCAGGACTTGCGGCGACGGTTCTTCCTTCACCATCT G-3′

### sgRNA sequences (including BsmBI Site)

sg*PTHLH* sense 5′-CACCGCATACGGGCAGCACGACGCG-3′

sg*PTHLH* antisense 5′-AAACCGCGTCGTGCTGCCCGTATG-3′

sg*HIF1α* sense 5’-CACCGTATGTGTGAATTACGTTGTG-3’

sg*HIF1α* antisense 5’-AAACCACAACGTAATTCACACATA-3’

sg*ARNT* sense 5’-CACCGTGGGGAACCTCACTTCGTGG-3’

sg*ARNT* antisense 5’-AAACCCACGAAGTGAGGTTCCCCA-3’

### Chemicals

PT2399 (obtained from Merck & Co., Inc., Rahway, NJ, USA) was reconstituted in sterile DMSO to achieve a stock concentration of 2 mM and added to media to achieve a final concentration of 2 μM. Biotin (Sigma-Aldrich; B4501-1G) was prepared to a stock solution of 200 mM in sterile water and added to media to achieve the desired final concentration of 12.5 µM. Doxycycline (Takara; 631311) was reconstituted in sterile water to achieve a stock concentration of 1 mg/mL and added to media to achieve a final concentration of 1 μg/mL. Zoledronic acid (Selleck Chemicals; S1314-100mg) was prepared as a stock solution of 1 mg/mL in PBS.

### Immunoblot analysis

Immunoblot analysis was performed as described elsewhere ^64^. The primary antibodies used included: Rabbit anti–Cyclin D1 (Cell Signaling Technologies; 2978), Rabbit anti-HIF2α (D6T8V) (Cell Signaling Technology; 29973S), Rabbit anti-NDRG1(Cell Signaling Technology; 5196S), Mouse anti-Vinculin (Sigma; V9131), Rabbit anti-V5-Tag (Cell Signaling Technology; 13202S), Mouse anti-b-Actin (Cell Signaling Technology; 3700S), anti-streptavidin-HRP (Cell Signaling Technology; 3999S), Rabbit anti-IGFBP3 (Cell Signaling Technology; 25864S), and Rabbit anti-MMP2 (D2O4T) (Cell Signaling Technology; 87809S).

### Lentivirus preparation and infection

Lentivirus preparation and infection were performed as described in ^64^. Briefly, lentiviruses were prepared by seeding 1.5 x 10^6^ 293T cells in a 60 mm plate, followed the next day by transfection with 1 μg of lentiviral vector DNA, 0.75 μg psPAX2, and 0.25 μg pMD2.G in 250 μL Opti-MEM with 6 μL TransIT-VirusGEN reagent (Mirus; MIR 6700). After a 15-minute incubation at room temperature, the mixture was added dropwise to 293T cells in 3 mL P/S-free DMEM with 10% FBS. The cells were incubated overnight, and then the medium was replaced with DMEM containing 30% FBS and 1% P/S. Virus-containing media were collected at 24 and 48 hours, pooled, centrifuged, filtered, and stored in 1.5 mL aliquots at -80°C. For infection, 2 x 10^6^ target cells were plated in 6-well plates with 1 mL of lentivirus, 10 μg/mL polybrene (Santacruz; sc-134220), and 1.5 mL media per well. Plates were gently shaken, centrifuged at 200 x g for 30 minutes at 30°C, then incubated overnight. The following day, cells were trypsinized, transferred to 10 cm plates, and cultured with the appropriate drug for selection of stable cell lines.

### BirA-ER labeling and streptavidin pulldown

For each condition to be tested, 4 x 10^6^ cells were seeded in 30 mL culture medium in 15 cm dishes. The next day, the cells were rinsed three times with Dulbecco’s Phosphate-Buffered Saline (DPBS) (Gibco; 14190144) to remove residual culture media. The cells were then treated with either DMSO, 2 µM PT2399, DMSO + 12.5 µM biotin, or PT2399 + 12.5 µM biotin in 18 mL DMEM supplemented with 1% Pen/Strep and 0.5% FBS, in duplicate. Forty eight hours later the media was collected, centrifuged at 50 x g for 10 minutes at 4°C and filtered through a 0.45 µm filter to remove cellular debris. The media was concentrated using Amicon Ultra-15 concentrators (Millipore; UFC900324) by two successive spins at 358 x g for 15 minutes at 4°C in an Eppendorf 5810R-15 amp version tabletop centrifuge, each time discarding the flow-through. To fully remove residual biotin, 15 mL DPBS was added to the concentrated media sample and centrifuged using the same centrifugation conditons, again discarding the flow-through. If more than 1 mL remained after the final spin, additional spins were conducted until the retained volume was reduced to ∼1 mL, which was then transferred to a 1.5 mL Eppendorf tube.

In parallel, cell lysates were also prepared to assess intracellular biotin labeling and protein expression. After removing media for the pulldown experiments, cell monolayers were rinsed twice with DPBS and detached from the culture surface with a cell lifter in 1 mL PBS. The cell suspension was then transferred to a 1.5 mL Eppendorf tube and pelleted in a benchtop centrifuge Eppendorf 5810R-15 amp version tabletop centrifuge at 50 x g for 5 minutes at 4°C. DPBS was aspirated and the cell pellet was lysed in 100 µL RIPA Buffer (Life Technologies; 89901) supplemented protease (Roche; 1836170) and phosphatase (Sigma Aldrich; 04906837001) inhibitors and rotated at 4°C 30 minutes. Lysates were cleared of insoluble debris by centrifugation at 17,000 x g for 15 minutes at 4°C in Eppendorf microcentrifuge and stored at -80°C or used immediately for Western blot analysis.

To prepare streptavidin magnatic beads (ThermoFisher; 65002), the beads were first resuspended by thorough vortexing until no beads adhered to the bottom of the bottle when inverted. 500 µL of resuspended beads were transferred to a 15 mL conical tube, adjusted to 10 mL with Tris-buffered saline with 0.1% Tween 20 (TBS-T), and spun at 358 x g in a Eppendorf 5810R-15 amp version tabletop centrifuge for 10 minutes at 4°. The supernatant was removed by aspiration with a 10 mL pipette and the beads were resuspended in 1 mL RIPA Buffer supplemented with protease and phosphatase inhibitors and transferred to 1.5 mL Eppendorf tubes. The tubes with the beads were transfered to a magnetic strip, washed twice with fresh 1 mL RIPA, and resuspended in 500 µL TBS-T. Beads were stored on ice until use.

For capture of biotinylated proteins, the volume of each concentrated media sample was brought up to ∼1 mL with DPBS. 30 µL of resuspended streptavidin beads were then added, ensuring thorough beads resuspension prior to addition. Samples were incubated overnight on an end over end rotator at 4°C. The next day, the tubes were quick-spun in an Eppendorf microcentrifuge to collect any liquid from the caps and then tranferred to a magnetic strip to pellet the beads. The supernatant was removed by aspiration with a P1000 pipette. The beads were washed twice with 1 mL RIPA Buffer containing protease and phosphatase inhibitors, followed by sequential washes with 1 mL 1 M KCl, 0.1 M Na_2_CO_3_ followed by 2 M Urea in 10 mM Tris-HCl pH 8. The beads were then washed twice with RIPA, twice with DPBS, and finally resuspended in 1 mL DPBS. The urea washes were performed in batches of no more than four samples, immediately followed by a RIPA wash to preserve sample integrity. A 35 µL aliquot of each bead suspension was removed for quality control, and the remainder was stored at -80°C for Mass Spectrometry analysis. For QC, the beads were pelleted by a quick spin in an Eppendorf microcentrifuge. The DPBS was carefully aspirated and the beads were resuspended in 30 µL of 1x Laemmli Sample buffer with 7.5 mM biotin to facilitate protein elution. For quality control experiments the beads were boiled at 95°C for 10 minutes, then either frozen at -80°C or used immediately for Western blot analysis. Otherwise the beads were used for on-bead digestion, as described below.

### On-bead trypsin digestion of biotinylated proteins

Peptides bound to streptavidin magnetic beads were washed four times with 200 µL of 50 mM Tris-HCl (pH=7.5) buffer. After removing the final wash, the beads were incubated twice at room temperature (RT) in 80 µL of the digestion buffer – 2 M Urea, 50 nM Tris-HCl, 1 mM DTT, and 0.4 µg trypsin – while shaking at 1000 rpm. The first incubation lasted 1 hour, followed by the second incubation of 30 minutes. After each incubation, the supernatant was collected and transferred into a separate tube. The beads were then washed twice with 60 µL of 2 M Urea/ 50 mM Tris-HCl buffer. The resulting washes were combined with the digestion supernatant. The pooled eluate of each sample was then spun down at 5000 x g for 30 sec to collect the supernatant. The samples were subsequently reduced with 4 mM DTT for 30 minutes at RT with shaking at 1000 rpm, followed by alkylation with 10 mM Iodoacetamide for 45 min in the dark at RT while shaking at 1000 rpm. Overnight digestion of the samples was performed by adding 0.5 µg of trypsin to each sample. The following morning, the samples were acidified with neat formic acid (FA) to the final concentration of 1% FA (pH<3).

Digested peptide samples were desalted using in-house packed C18 (3M) StageTips. C18 StageTips were conditioned sequentially with 100 µL of 100% methanol (MeOH), 100 µL of 50% (vol/vol) acetonitrile (MeCN) with 0.1% (vol/vol) FA, and two washes of 100 µL of 0.1% (vol/vol) FA. Acidified peptides were loaded onto the C18 StageTips and washed twice with 100 µL of 0.1% FA. The peptides were then eluted from the C18 resin using 50 µL of 50% MeCN/ 0.1% FA. The desalted peptide samples were snap-frozen and vacuum-centrifuged until completely dry.

### TMT labeling and fractionation of peptides for analysis by LC-MS/MS

Desalted peptides were labeled with TMT16 reagents (Thermo Fisher Scientific). Each peptide sample was resuspended in 80 μL of 50 mM HEPES and labeled with 20 µL of the 25 µg/µL TMT reagents in MeCN. The samples were then incubated at RT for 1 hour while shaking at 1000 rpm. To quench the TMT-labeling reaction, 4 μL of 5% hydroxylamine was added to each sample, followed by a 15-minute incubation at RT with shaking. TMT-labeled samples were combined and vacuum-centrifuged to dry. The samples were then reconstituted in 200 μL of 0.1% FA and desalted on a C18 StageTip using the previously described protocol. The desalted TMT-labeled combined sample was then dried to completion.

The combined TMT-labeled peptide sample was fractionated by basic reverse phase (bRP) fractionation using an in-house packed SDB-RPS (3M) StageTip. A StageTip containing three plugs of SDB material was prepared and conditioned with 100 μL of 100% MeOH, 100 μL of 50% MeCN/0.1% FA, and 2x with 100 μL of 0.1% FA. The sample was resuspended in 200 µL 0.1% FA (pH < 3) and loaded onto the conditioned StageTip and eluted in a series of buffers with increasing MeCN concentrations. Six fractions were collected in 20 mM ammonium formate (5%, 10%, 15%, 20%, 25%, and 45% MeCN), dried to completion and analyzed by LC-MS/MS

### Liquid chromatography tandem mass spectrometry

All peptide samples were separated and analyzed on an online liquid chromatography tandem mass spectrometry (LC-MS/MS) system, consisting of a Vanquish Neo UPHLC (Thermo Fisher Scientific) coupled to an Orbitrap Exploris 480 (Thermo Fisher Scientific). All peptide fractions were reconstituted in 9 µL of 3% MeCN/ 0.1% FA. 4 µL of each fraction was injected onto a microcapillary column (Picofrit with 10 µm tip opening/ 75 µm diameter, New Objective, PF360-75-10-N-5), packed in-house with 30 cm of C18 silica material (1.5 µm ReproSil-Pur C18-AQ medium, Dr. Maisch GmbH, r119.aq) and heated to 50 °C using column heater sleeves (PhoenixST). Peptides were eluted into the Orbitrap Exploris 480 at a flow rate of 200 nL/min. The bRP fractions were run on a 154 min-method. Solvent A comprised 3% acetonitrile/ 0.1% FA. Solvent B comprised 90% acetonitrile/ 0.1% FA. The LC-MS/MS method used the following gradient profile: (min: %B) 0:2; 1:6; 122:35; 130:60; 133:90; 143:90; 144:50; 154:50 (the last two steps at 500 nl/min flow rate).

Mass spectrometry was conducted using a data-dependent acquisition mode, MS1 spectra were measured with a resolution of 60,000, a normalized AGC target of 100%, and a mass range from 350 to 1800 m/z. MS2 spectra were acquired for the top 20 most abundant ions per cycle at a resolution of 45,000, an AGC target of 50%, an isolation window of 0.7 m/z and a normalized collision energy of 32. The dynamic exclusion time was set to 20 s.

### Analysis of mass spectrometry data

Mass spectrometry data was processed using Spectrum Mill (proteomics.broadinstitute.org). Spectra within a precursor mass range of 600-6000 Da with a minimum MS1 signal-to-noise ratio of 25 were retained. Additionally, MS1 spectra within a retention time range of +/-45 s, or within a precursor m/z tolerance of +/-1.4 m/z were merged. MS/MS searching was performed against a human Uniprot database. For searching, fixed modifications were TMT16-Full-Lys modification and carbamidomethylation on cysteine. Variable modifications included acetylation of the protein N-terminus, oxidation of methionine and cyclization to pyroglutamic acid. Digestion parameters were set to “trypsin allow P” with an allowance of 4 missed cleavages. The matching tolerances were set with a minimum matched peak intensity of 30%, precursor and product mass tolerance of +/-20 ppm.

Peptide spectrum matches (PSMs) were validated with a maximum false discovery rate (FDR) threshold of 1.2% for precursor charges ranging from +2 to +6. A target protein score of 9 was applied during protein polishing auto-validation to further filter PSMs. TMT16 reporter ion intensities were corrected for isotopic impurities using the afRICA correction method in the Spectrum Mill protein/ peptide summary module, which utilizes determinant calculations according to Cramer’s Rule. Protein quantification and statistical analysis were performed using the Proteomics Toolset for Integrative Data Analysis (Protigy, v1.0.7, Broad Institute, https://github.com/broadinstitute/protigy). Differential protein expression was evaluated using moderated t-tests, with 2 sided P-values calculated to assess significance.

### Quantitative Real-Time PCR (qRT-PCR)

Total RNA was extracted from cells using the RNeasy Mini Kit (Qiagen; 74106) according to the manufacturer’s instructions. Complementary DNA (cDNA) was synthesized from 1 μg of total RNA using the iScript RT Supermix for RT-qPCR (Bio-Rad Laboratories; 1708841) following the manufacturer’s protocol. The synthesized cDNA was diluted 1:10 with nuclease-free water prior to use. qRT-PCR was performed on a LightCycler 480 II system (Roche Diagnostics) using the LightCycler 480 SYBR Green I Master Mix (Roche Diagnostics; 04887352001). Each reaction contained 1 μL of primer mix, 2 μL of diluted cDNA (1:10), 3 μL of nuclease-free water, and 5 μL of SYBR Green Master Mix, in a total volume of 10 μL. All reactions were run in triplicate, and relative gene expression levels were calculated using the ΔΔCt method, with target gene expression normalized to housekeeping controls.

### qRT-PCR primers

*PTHLH* Forward 5′- GAACTGGCTCTGCCTGGTTAGA -3′

*PTHLH* Reverse 5′- GTCCTTGGAAGGTCTCTGCTGA-3′

*NDRG1* Forward 5′- CTCCTGCAAGAGTTTGATGTCC -3′

*NDRG1* Reverse 5′- TCATGCCGATGTCATGGTAGG-3′

*IGFBP3* Forward 5′- CGCTACAAAGTTGACTACGAGTC-3′

*IGFBP3* Reverse 5′- GTCTTCCATTTCTCTACGGCAGG-3′

*BNIP3* Forward 5′- TCCTGGGTAGAACTGCACTTC-3′

*BNIP3* Reverse 5′- GCTGGGCATCCAACAGTATTT-3′

*IGFBP4* Forward 5′-ACCCACGAGGACCTCTACATCA -3′

*IGFBP4* Reverse 5′- CACACCAGCACTTGCCACGCT -3′

*IGFBP6* Forward 5′- CACAGGATGTGAACCGCAGAGA-3′

*IGFBP6* Reverse 5′- CACTGAGTCCAGATGTCTACGG-3′

*IGF1* Forward 5′- GCTCTTCAGTTCGTGTGTGGA -3′

*IGF1* Reverse 5′- GCCTCCTTAGATCACAGCTCC-3′

*SFRP4* Forward 5′- ACGAGCTGCCTGTCTATGAC-3′

*SFRP4* Reverse 5′- TGTCTGGTGTGATGTCTATCCAC-3′

*IL6* Forward 5′- AGACAGCCACTCACCTCTTCAG-3′

*IL6* Reverse 5′- TTCTGCCAGTGCCTCTTTGCTG-3′

*CXCL8* Forward 5′- GAGAGTGATTGAGAGTGGACCAC-3′

*CXCL8* Reverse 5′- CACAACCCTCTGCACCCAGTTT-3′

*GDF15* Forward 5′- CAACCAGAGCTGGGAAGATTCG-3′

*GDF15* Reverse 5′- CCCGAGAGATACGCAGGTGCA-3′

*EGLN3* Forward 5′- TCCTGCGGATATTTCCAGAGG-3′

*EGLN3* Reverse 5′- GGTTCCTACGATCTGACCAGAA-3′

*VEGFA* Forward 5′- AGGGCAGAATCATCACGAAGT -3′

*VEGFA* Reverse 5′- AGGGTCTCGATTGGATGGCA -3′

*PTHLH*-V5-ORF Forward 5′- AACGTCGCTGGAGCTCG -3′

*PTHLH*-V5-ORF Reverse 5′- GTGGGTTTGGGATTGGCTTTCC -3′

*UBC* Forward 5′- CTGGAAGATGGTCGTACCCTG -3′

*UBC* Reverse 5′- GGTCTTGCCAGTGAGTGTCT -3′

*ACTB* Forward 5′- CATGTACGTTGCTATCCAGGC-3′

*ACTB* Reverse 5′- CTCCTTAATGTCACGCACGAT -3′

### Colorimetric calcium assay

Serum calcium concentrations were measured using a colorimetric calcium assay kit (ab102505; Abcam) according to the manufacturer’s instructions. Briefly, 10 µL of each serum sample was added to the well of a 96-well plate and brought to a total volume of 50 µL with deionized water. 90 µL of chromogenic reagent and 60 µL of calcium assay buffer was then added to each well, followed by incubation at room temperature in the dark for 5 –10 minutes. Absorbance was measured at 575 nm using a microplate reader. A calcium standard curve was generated with a series of known calcium concentrations, and sample concentrations were calculated based on this curve.

### PTHrP and GDF15 ELISA Assay

We measured the PTHrP 1-34 active peptide in cell culture media using the Parathyroid Hormone-Related Protein (PTHrP) (1-34) EIA Kit, extraction-free (Phoenix Pharmaceuticals, INC; EK-056-04) and GDF15 using the Human GDF-15 ELISA Kit – Quantikine (R&D Systems; DGD150) according to the manufacturer’s instructions. For each assay, 0.3 x 10^6^ cells were seeded per well in a 6-well plate. The following day, the media was replaced with 2 mL DMEM or RPMI containing 1% Pen/Strep and 0.5% FBS, and, where indicated, the indicated treatment. After 48 hours, media were collected and centrifuged at 400 x g for 2 minutes to pellet any debris, and the supernatant was saved. Simultaneously, cells were lysed in 150 µL of lysis buffer, and total protein concentration was measured. 50 µL of conditioned media was used to measure PTHrP (ng/ mL) or GDF15 (pg/ mL) concentration in unit, which was subsequently normalized to the total cellular protein (μg), where total cellular protein = total cellular protein concentration (μg/ mL) x 0.15 mL.

### Knock-In FLAG-HA Tag at the Endogenous HIF2α Locus

The generation of 3×FLAG-HA-HIF2α knock-in (KI) cells was performed following the protocol published by Jiang et al. ^65^, summarized here. SPRI magnetic beads (GE Healthcare; 65152105050250) were used for purification of the HDR template synthesized to introduce a 3×FLAG-HA tag at the N-terminus of HIF2α. Cas9-RNP complexes were assembled using Alt-R Cas9 Nuclease V3 (stock solution: 62 mM, IDT 1081059), synthetic sgRNA (IDT), and Alt-R Cas9 Electroporation Enhancer (IDT). OSRC-2 cells (2 x 10⁵) were electroporated with Cas9-RNPs and HDR template using the 4D-Nucleofector X unit (Lonza) with condition EN138. Post-electroporation, cells were cultured in media supplemented with Alt-R HDR Enhancer V2 (IDT) and refreshed after 16 hours. Knock-in efficiency was confirmed by amplicon sequencing or immunoblot analysis 3–4 days later. Single-cell clones were generated for subsequent ChIP-seq assays.

### Chromatin Immunoprecipitation (ChIP)

ChIP was performed as described in Jiang et al. ^65^. Briefly, 1×10⁷ OSRC-2 cells were crosslinked using 1% formaldehyde, which was then quenched with glycine. Crosslinked cells were lysed in SDS lysis buffer (1% SDS, 10 mM EDTA, 50 mM Tris-HCl, pH 8.0) supplemented with protease inhibitors and 5mM sodium butyrate. Chromatin was fragmented by sonication (Covaris E220) to an average size of ∼200-500 bp. For ChIP-seq, 40 µg of chromatin was immunoprecipitated using 5 µg FLAG antibody (Sigma-Aldrich; F1804). Antibody-bead conjugates were prepared using Protein G magnetic beads (Life Technologies; 10004D) and washed extensively. Bead-bound chromatin was washed sequentially with RIPA 0, RIPA 0.3, and LiCl buffers before elution in SDS elution buffer. DNA was reverse crosslinked at 65°C and purified using the MinElute PCR Purification Kit (Qiagen; 28004). ChIP-seq libraries were prepared using the Swift DNA Library Prep Kit and sequenced on an Illumina NextSeq 500 platform. Peak calling and differential binding analyses were performed using MACS2 and DEseq2 within the CoBRA pipeline as described by Jiang et al. ^65^.

### RNA-seq Sample Library Preparation and Sequencing

For 72 h PT2399 treatment and sgHIF2 experiments, RNA-seq was done as described in Jiang et al.^65^. In brief, ribosomal RNA (rRNA)-depleted libraries were prepared from 100 ng of total RNA using the KAPA RNA HyperPrep Kit with RiboErase (Roche, #08098131702). RNA was fragmented at 94°C for 8 minutes, followed by first- and second-strand cDNA synthesis. cDNA fragments were end-repaired, adenylated at the 3′ ends, and ligated to universal adapters. Indexed libraries were enriched by 14 cycles of PCR using a Beckman Coulter Biomek i7 Automated Workstation. Libraries were quantified using a Qubit 4 Fluorometer and Agilent TapeStation 4200, pooled in an equimolar ratio, and shallowly sequenced on an Illumina MiSeq to assess quality. Final sequencing was conducted with paired-end 150 bp reads on an Illumina NovaSeq 6000 at the Dana-Farber Cancer Institute Molecular Biology Core Facilities.

For 24 h and 48 h PT2399 treatment experiments, cells were seeded at a density of 1 x 10⁶ cells per 100 mm tissue culture dish (Corning, 353003) and, in triplicate, treated with either DMSO or 2 µM PT2399 for 24 or 48 hours. Following treatment, the cells were rinsed twice with ice-cold Dulbecco’s Phosphate Buffered Saline (DPBS) (Gibco, 14190094) and scraped into 1 mL of ice-cold DPBS. The cell suspension was centrifuged at 1200 x g for 5 minutes at 4°C to pellet the cells. The pellets were then flash-frozen in liquid nitrogen and stored at –80°C until processing.

For RNA extraction, the Qiagen RNeasy Mini Kit (Qiagen, 74106) was used. RNA quantity and quality were measured on a NanoDrop spectrophotometer (Thermo Scientific, ND-8000-GL). The samples were subsequently sent to GENEWIZ for library preparation and sequencing. Prior to library construction, normalization was achieved by adding the ERCC RNA Spike-In Mix kit (ThermoFisher Scientific, 4456740) according to the manufacturer’s protocol.

Library construction began with the enrichment of mRNA using Oligo(dT) beads. The enriched mRNA was then fragmented by incubation with NEBNext First Strand Synthesis Reaction Buffer at 94°C for 15 minutes. First and second strand cDNA synthesis were performed next, followed by end repair and 3’ adenylation of the cDNA fragments. Universal adapters were ligated, and indexed PCR amplification (with a limited number of cycles) was carried out to enrich the library. Library quality was verified using an Agilent TapeStation (Agilent Technologies, G2991BA) and quantified with both a Qubit 2.0 Fluorometer and quantitative PCR (KAPA Biosystems, KK4824).

Clusters were generated on lanes of a single flow cell, which was then loaded onto an Illumina HiSeq instrument (4000 or equivalent) per the manufacturer’s instructions. Sequencing was performed using a 2 x 150 bp paired-end configuration. Image analysis and base calling were executed with HiSeq Control Software (HCS), and the raw .bcl files were converted to FASTQ files and demultiplexed using Illumina’s bcl2fastq 2.17 software, allowing for one mismatch in index sequence identification.

The paired-end FASTQ files were aligned to the hg38 genome using HISAT2. Gene-level read counts were obtained with featureCounts and normalized using DESeq2, which modeled the average gene expression in each sample with a negative binomial distribution. Differential expression analysis between PT2399-treated (24 or 48 hours) and DMSO-treated samples was performed using a Wald test provided by DESeq2, and corresponding plots were generated to illustrate these differences.

### PRO-seq Sample Preparation and Analysis

Pro-seq was done as described in Jiang et al. ^65^. Briefly, cells were harvested, permeabilized, and flash frozen in freezing buffer. Nascent RNA was labeled using biotin-11-NTPs in nuclear run-on assays and purified using streptavidin beads. Following fragmentation and adapter ligation, reverse transcription was performed to generate cDNA, which was amplified to construct sequencing libraries. Libraries were sequenced using the Illumina NovaSeq platform, and the data were analyzed with scripts developed by the AdelmanLab. Transcription start sites were annotated using Ensembl GTF files, and browser snapshots were generated with IGV for visualization.

### Polysome-Seq Sample Preparation

Polysome-Seq was done as described in Jiang et al. ^65^. Briefly, 1 x 10⁷–2 x 10⁷ OSRC-2 cells were washed with ice-cold PBS and lysed on ice using polysome extraction buffer (25 mM HEPES pH 7.5, 5 mM MgCl₂, 100 mM KCl, 2 mM DTT, 1% Triton X-100, 0.1 mg/mL cycloheximide, 0.05 U/μL RNase inhibitor, and 1x EDTA-free protease inhibitor cocktail). Lysates were homogenized with a glass Dounce homogenizer and cleared by centrifugation (10,000 x g, 5 min, 4°C). The supernatant was loaded onto 10–50% sucrose gradients and centrifuged (40,000 rpm, 3 h, 4°C, SW41Ti rotor). Gradients were fractionated, and polysome-containing fractions (all heavier than the monosome peak) were pooled for RNA purification using TRIzol LS, followed by sequential isopropanol and ethanol precipitations. RNA quality was verified (RIN > 9) using the Agilent RNA TapeStation, and sequencing libraries were prepared using the SMARTer Stranded Total RNA Sample Prep Kit - HI Mammalian. Final libraries were quantified with Qubit HS DNA assays and an Agilent TapeStation, pooled with unique dual indexes, and sequenced on an Illumina NovaSeq 6000.

### Differential gene expression analysis for RNA-seq and Polysome-seq

Sequenced reads were aligned to the UCSC hg38 reference genome assembly and gene counts were quantified using STAR (v2.7.3a) ^66^ and Salmon ^67^. Differential gene expression testing was performed by DESeq2 (v1.22.1) ^68^. RNAseq analysis was performed using the VIPER snakemake pipeline ^69^.

### Mouse xenograft models

All experimental procedures involving orthotopic and subcutaneous xenografts were approved by the Dana-Farber Cancer Institute or the University of Massachusetts Chan Medical School Institutional Animal Care and Use Committees. All in vivo experiments were performed using female mice. Female mice were selected based on prior experience with the OSRC-2 xenograft model, which showed consistent tumor engraftment, and because they are generally easier to handle, reducing stress-related variability. Female Ncr nude mice (NCRNU-F sp/sp CrTac:NCr-*Foxn*1^nu^) were obtained from Taconic and Female NOD Cg-*Prkdc*^scid^ *Il2rg*t^m1Wjl^/SzJ (NGS) mice were obtained from The Jackson Laboratory. To establish orthotopic xenografts, 1 x 10^6^ 786-O-Fluc cells or 0.5 x 10^6^ OSRC-2-Fluc cells were injected into the parenchyma of the left kidney of nude mice, as described by us previously ^26^. For subcutaneous xenografts, 5 x 10^6^ OSRC-2-Fluc cells were injected into the right flank of nude mice or 6 x 10^6^ RXF393-Fluc cells were injected subcutaneously into both the right and left flanks of nude mice or NSG mice. For the data in Fig. 1 and Extended Data Fig. 6, tumors were allowed to form, and mice body weights were monitored. When a mouse lost > 10% of its body weight, blood was drawn to collect serum for calcium measurement. Mice were then randomized to receive either 45 mg/kg PT2399 (formulated in 10% ethanol, 30% PEG400, and 60% water containing 0.5% methylcellulose and 0.5% Tween 80) or vehicle alone, administered daily by oral gavage for 6 days. Body weight was monitored daily. For the zoledronic acid experiments for Extended Data Fig. 4h–n, OSRC-2 tumor– bearing mice were randomized into three treatment arms: vehicle, PT2399 (45 mg/kg daily oral gavage), or zoledronic acid (ZOL; 120 μg/kg intraperitoneally every other day). For the data in Fig. S1, PT2399 began 4 weeks after tumor cell implantation and was dosed at 30 mg/kg daily by oral gavage. In experiments with the DOX inducible experiments, mice were fed a normal chow diet (-DOX) or, where indicated, a Doxycycline 2000 PPM Green diet (TestDiet; 1811824). At the study endpoint, mice were euthanized using CO₂. Tumors were then harvested and weighed. A portion of each tumor was frozen at -80°C, while the remainder was fixed in 10% paraformaldehyde for histological analysis.

For the radiological and histological characterization of the cachexia in OSRC-2 cells model, NCr nude female mice were purchased from Taconic biosciences (NCRNU-F sp/sp CrTac:NCr-*Foxn*1^nu^) and were allowed to acclimatize for a week with *ad libitum* access to food and water. Mice were implanted with 5 x 10^6^ OSRC-2 cells (or sham controls for non-tumor bearing) subcutaneously and weighed every other day.

Tumor-bearing mice were randomized into vehicle or PT2399-treatment (30mg/kg) arms and received one dose daily by oral gavage for a total of 17 days. Body scans were performed on an Echo Medical Systems 1H-MRS by the UMass Metabolic Disease Research Center (MDRC) to measure the fat and lean masses of each animal right before the first dose of drug and 17 days into treatment. Mice were housed in TSE metabolic cages in the MDRC facility, where daily food intake was monitored over a three day period. After the last treatment, mice were euthanized and tumor, iWAT, eWAT, BAT, gastrocnemius, and quadriceps tissues were collected, weighed, and processed for downstream analysis. For H&E (Hematoxylin and Eosin) and IF (Immunofluorescent) staining, dissected tissue was immediately fixed in Zinc Formalin, processed, embedded in paraffin, sectioned, and mounted. Immunofluorescent sections were deparaffinized and rehydrated, followed by antigen retrieval and overnight incubation with Ucp1 (Sigma CAT #U6382) primary antibody at 4°C at 1:1000. Images were acquired using an EVOS M5000 fluorescent microscope and quantification of percent area was performed using ImageJ software.

### *PTHLH* mRNA expression and copy number analysis

The gene expression files for each TCGA sample (“*.rna_seq.augmented_star_gene_counts.tsv”) were downloaded from the GDC Data Portal, and TPM values were extracted. Only data from primary tumor samples were included in the analysis. Copy number variation (CNV) data for PTHLH were retrieved from the “PanCancer Atlas” dataset in cBioPortal. Copy numbers corresponding to the PTHLH gene were classified as follows: cn = 2 (Amplification), cn = 1 (Gain), cn = 0 (Diploid), cn = -1 (Heterozygous deletion), and cn = -2 (Homozygous deletion). Data were restricted to primary tumor samples. To analyze the relationship between copy number and gene expression levels for genes on chromosome 12p in KIRC, samples were grouped based on CNV status (Amplification or Gain vs. no copy number change), and the mean log2 fold change in gene expression was calculated. Statistical differences between groups were assessed using the unpaired Wilcoxon rank sum test. Additionally, relative expression levels normalized to *ACTB* were calculated for samples with Amplification or Gain, and the mean values were reported.

### Clinical data for belzutifan, ICI, and VEGF TKI

Plasma samples were obtained for patients with advanced ccRCC treated at the Dana-Farber Cancer Institute under protocol #01-130. Matched pre-and post-treatment samples (250 µL each) were collected from patients receiving either the HIF2 inhibitor (belzutifan), immune checkpoint inhibitors or VEGF tyrosine kinase inhibitors. Plasma PTHrP levels were measured using an Immunochemiluminometric Assay at the Mayo Clinic (Test Id: PTHRP). Clinical data, including baseline characteristics, body weight, and albumin-corrected calcium levels at the time therapy was initiated, 1 month, and 3 months thereafter were retrospectively collected. Variations in PTHrP levels, corrected calcium, and body mass index (BMI) were evaluated using paired Wilcoxon signed-rank tests. Both male and female patients with RCC were included in this study and were reported in Extended Data Table 1. Sex was determined from medical records as sex assigned at birth. Analyses were performed disaggregated by sex where relevant and included in the source data.

### Clinical data for NKT2152

Plasma samples (in K2 EDTA tubes) were collected from 2021 to 2024 in the dose escalation part of a phase I study of NKT2152, a HIF2 Inhibitor, in patients with advanced or metastatic ccRCC (NCT05119335). To be eligible for this study patients had to be aged 18 years or older and with locally advanced or metastatic ccRCC and to have exhausted available standard therapy as determined by the investigator. 60 subjects were enrolled in the dose escalation part, among which 45 subjects had plasma samples for the PTHrP assay and data analysis. PTHrP level was measured at Mayo Clinic with a Immunochemiluminometric Assay (Test Id: PTHRP). Sample shipping was at -20°C with dry ice; sample storage was at -70°C. There was a freeze/thaw cycle for the transfer of the plasma samples stored in a 2.0 mL Cryovial tube to a 5.0 mL test tube before the test. Data Analysis: For PTHrP results < 0.4 pmol/L, 0.2 (1/2 of LOQ) was used for plots. Windowing algorithm and Last Observation Carried Forward (LOCF) were applied for plotting of longitudinal body weight and Calcium level changes. Clinical data cutoff date was on June 16, 2024. Both male and female patients with RCC were included in this study. Sex was determined from medical records as sex assigned at birth and were reported in Extended Data Table 1. Analyses were performed disaggregated by sex where relevant and included in the source data.

### Statistical Analysis

Statistical analyses were performed using GraphPad Prism and R. Data are presented as mean ± SEM. Normality was visually assessed, and appropriate statistical tests were selected accordingly.

For comparisons between two groups, unpaired two-tailed Student’s t-tests or unpaired Wilcoxon rank sum tests were used, depending on data distribution. For paired data comparisons, the paired Wilcoxon signed-rank test was applied. For multiple group comparisons, two-way ANOVA was performed. Kaplan-Meier survival analyses were assessed using the log-rank test.

Scatterplots were generated to visualize the relationship between specific tissue mass measurements and pre-treatment weights. Analysis of Covariance (ANCOVA) was utilized to address the relationship of specific mass measurements and treatment group to specifically compare vehicle to PT2399, when the similar slope assumption was met. All models were adjusted for pre-treatment body weight. A heteroskedasticity-consistent variance-covariance matrix was employed to calculate robust estimates.

For omics data analysis, RNA-seq differential expression analysis was performed using DESeq2, with statistical significance determined by the Wald test. ChIP-seq peak calling was conducted using MACS2, and proteomics data were normalized using Spectrum Mill.

All statistical details, including the number of samples (n), statistical tests used, and significance thresholds, are provided in the figure legends and in the relevant Methods sections. The n represents biological replicates, not technical replicates.

Mice were randomly assigned to treatment groups, and blinding was applied for histological assessments. All experiments were replicated at least twice. Statistical significance was set at 2 sided p ≤ 0.05, with exact p-values reported where applicable.

## Data availability

There is no restriction on experimental data availability for this study. The original mass spectra and the protein sequence databases used for searches have been deposited in the public proteomics repository MassIVE (http://massive.ucsd.edu) and are accessible at ftp://MSV000097181@massive.ucsd.edu when providing the dataset password: cachexia. If requested, also provide the username: MSV000097181. These datasets will be made public upon acceptance of the manuscript. The processed dataset and the data tables from Fig. 2, the uncropped western blot scans, and the sex-stratified analysis of Belzutifan and NKT2152 clinical data in RCC patients for Fig. 6 and Extended Data Fig. 8 were uploaded on Zenodo and will be publicly available after publication of the manuscript. The reviewers can access the data privately at link: https://zenodo.org/records/14953955?token=eyJhbGciOiJIUzUxMiJ9.eyJpZCI6IjEyMmViOTA5LWMzZmYtNDg5MC1hY2YzLTAzODczMjg1ZmYwNCIsImRhdGEiOnt9LCJyYW5kb20iOiJjNGQ0NTg0YWM3ODZmNDkyZmE5NWY5M2Q0MTFmMGVhMyJ9.irkwgZwS9dCEqjqVQVuMfsDooyH_qH5X-WmNxqWtJYwYWMGJ9zQyXQUIX76LXD7k6JrHxJogvnzWPGKe0t4y5A. All raw and processed dataset files from Fig. 3 were uploaded on Gene Expression Omnibus (GEO) and will be publicly available after publication of the manuscript. The RNA-Seq (24h and 48h PT2399 treatment), ChIP-Seq, and PRO-Seq files are publicly available and can be downloaded by using GEO accession number GSE277046. The reviewers can access the data at the link : https://www.ncbi.nlm.nih.gov/geo/query/acc.cgi?acc=GSE277046. The RNA-Seq (72h PT2399 treatment and sgHIF2α) files can be downloaded by using GEO accession number GSE289579. The reviewers can access the data privately at the link https://www.ncbi.nlm.nih.gov/geo/query/acc.cgi?acc=GSE289579 by using the private token “czkzsokgnpetrwv”. The Polysome-Seq files can be downloaded by using GEO accession number GSE289581. The reviewers can access the data privately at the link https://www.ncbi.nlm.nih.gov/geo/query/acc.cgi?acc=GSE289581 by using the private token “cxexggcyhfotvuh”.

## Code availability

The code used to analyze ChIP-seq, RNA-seq (24h and 48h of PT2399 treatment), and PRO-seq experiments’ datasets in this manuscript was deposited at kaelinlabdfci GitHub (https://github.com/kaelinlabdfci/HIF2a_CCND1_ccRCC). The code used to analyze the TCGA data was deposited at kaelinlabdfci GitHub (https://github.com/kaelinlabdfci/HIF2-RCC-Cachexia) and will be made publicly available after the publication of the manuscript.

## Acknowledgments

The authors thank Merck & Co., Inc., Rahway, NJ, USA for providing PT2399. W.G.K. is a Howard Hughes Medical Institute Investigator and is supported by NIH grant 2R35CA210068, and by the Dana-Farber/Harvard Cancer Center (DF/HCC) Kidney Cancer Specialized Program of Research Excellence (P50-CA101942-12). N.H.S. is supported by the Hope Funds for Cancer Research Sakonnet Family Fellowship and the Career Enhancement Program Award, Dana-Farber/Harvard Cancer Center Kidney Cancer Specialized Program of Research Excellence (P50-CA101942-12). We sincerely thank Nicholas J. Kramer and L. Stirling Churchman (Harvard Medical School) for assistance with the Polysome-seq experiments, Zachary T. Herbert and Molecular Biology Core Facilities at Dana-Farber Cancer Institute for help with performing and analyzing the RNA-seq and Polysome-seq experiments, the Gelb center and Mary Lee at DFCI for providing the patients’ plasma samples, the staff of the Dana-Farber Cancer Institute Animal Resource Facility for the care of research animals that enabled this study, and the W.G.K. laboratory members for sharing reagents and for their valuable discussions. The metabolic phenotyping of OSRC-2 tumor bearing animals was performed at the Metabolic Disease Research Center of the University of Massachusetts Chan Medical School and supported by NIH grants (5R01-DK133772 and 5R01-AG085308 to Jason K. Kim). We thank Wanling Xie and Morgan Paul and for their assistance with statistical power calculations and data analysis. We thank Yamini Ogoti and Lauren Tauer for their assistance in performing the murine body scans and quantification of fat and lean masses. T.K.C is supported by DF/HCC Kidney SPORE (2P50CA101942-16) DF/HCC Core Grant 5P30CA006516-56, the Kohlberg Chair at Harvard Medical School, the Trust Family, Michael Brigham, Pan Mass Challenge, Hinda and Arthur Marcus Fund, and Loker Pinard Funds for Kidney Cancer Research at DFCI. This work was supported in part by grants P01CA206978 to S.A.C from the NIH, and U24CA270823, U01CA271402 to S.A.C, from National Cancer Institute (NCI) Clinical Proteomic Tumor Analysis Consortium program, as well as a grant from the Dr. Miriam and Sheldon G. Adelson Medical Research Foundation to N.D.U. and S.A.C. This article is subject to HHMI’s Open Access to Publications policy. HHMI lab heads have previously granted a nonexclusive CC BY 4.0 license to the public and a sublicensable license to HHMI in their research articles. Pursuant to those licenses, the author-accepted manuscript of this article can be made freely available under a CC BY 4.0 license immediately upon publication.

## Author contributions

M.A.-R., L.A.S., and W.G.K. conceived the study. M.A-R. designed and performed the necessity and sufficiency experiments that validated PTHrP as the cause of cachexia in ccRCC models, including making the necessary cell lines and performing the in vivo treatments with PT2399. He also analyzed the data and generated all the figures. N.B. and J.R.P. performed the metabolic modeling for the mice with cachexia. L.A.S. and S.M.V. made the initial observation that PT2399 reversed the cachexia caused by OSRC-2 cells and conducted the BirA-ER labeling experiments. H.W., C.X., S.A.C., and N.D.U. performed the LC-MS/MS experiments and analyzed the mass spectrometry data. N.H.S. conducted ChIP-seq, RNA-seq (24h and 48h of PT2399 treatment), and PRO-seq experiments. Q.J. performed RNA-seq (72h PT2399 treatment and sgHIF2) and polysome-seq experiments. E.S. and T.K.C. analyzed clinical data from patient samples treated with belzutifan, ICIs, and TKIs. J.L., H.W., Z.L., and W.S. analyzed clinical data from patient samples treated with NKT2152. K.E. analyzed TCGA data. M.A.-R. and W.G.K. wrote the manuscript. W.G.K. supervised the study.

