## Extended Data Figures 1-8 for "HIF2-driven PTHrP Causes Cachexia and Hypercalcemia in Kidney Cancer: Treatment with HIF2 Inhibitors"

Extended Data Fig. 1

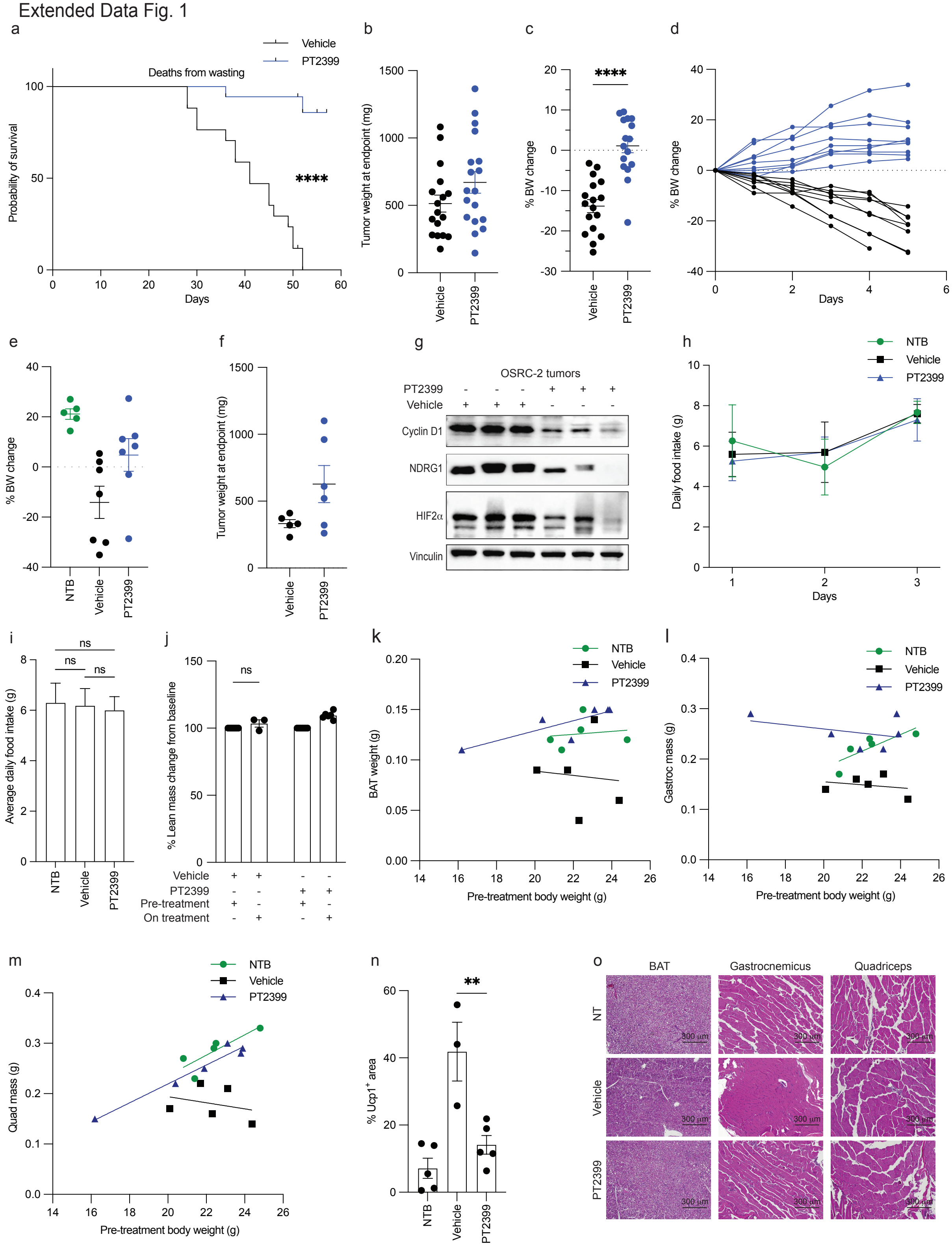

**Extended Data Figure 1. Supplemental data supporting the efficacy of PT2399 in treating cachexia induced by OSRC-2 xenografts.** (a) Kaplan-Meier curve showing the probability of survival for OSRC-2 orthotopically implanted tumor-bearing mice treated with PT2399 (blue, n = 18) or vehicle (black, n = 17). Deaths were attributed to wasting. Statistical analysis was performed using the log-rank test (****$P \leq 0.0001$). (b-c) (b) Tumor weight (c), body weight changes of mice in (a). (d) Individual trajectories of BW changes corresponding to Fig. 1b. (e-j) (e) Body weight change, (f) tumor weight, (g) immunoblot analysis of representative OSRC-2 tumor lysates, (h) daily food intake in grams over 3 days, (i) average daily food intake in grams, (j) percent fat mass change from baseline, measured by MRS body scan, in mice treated with PT2399 or vehicle. (k–o) (k) Brown adipose tissue (BAT) mass versus pre-treatment body weight. (l) Gastrocnemius (Gastroc) muscle mass versus pre-treatment body weight. (m) Quadriceps (Quad) muscle mass versus pre-treatment body weight. Each dot represents an individual mouse; regression lines are shown for group comparisons. ANCOVA adjusting for pre-treatment body weight revealed significantly increased BAT and Gastroc mass in PT2399-treated tumor-bearing mice compared to vehicle-treated controls (BAT: p = 0.023, 95% CI: 0.01–0.10 g; Gastroc: p = 0.001, 95% CI: 0.06–0.15 g). An interaction between treatment and pre-treatment body weight was detected for quadriceps mass, precluding ANCOVA analysis for that outcome, (n) quantification of Ucp1 positive area corresponding to Figure 1j, (o) Hematoxylin and eosin (H&E) staining of BAT, gastrocnemius muscle, and quadriceps muscle sections from tumor-bearing mice treated with vehicle or PT2399 compared to  non-tumor bearing (NTB) mice. Data are presented as mean ± SEM. Statistical significance was assessed using unpaired two-tailed t-tests. P values are indicated as follows: $P \leq 0.05$ (*), $P \leq 0.005$ (**), $P \leq 0.005$ (***), and $P \leq 0.0005$ (****).

Extended Data Fig. 2

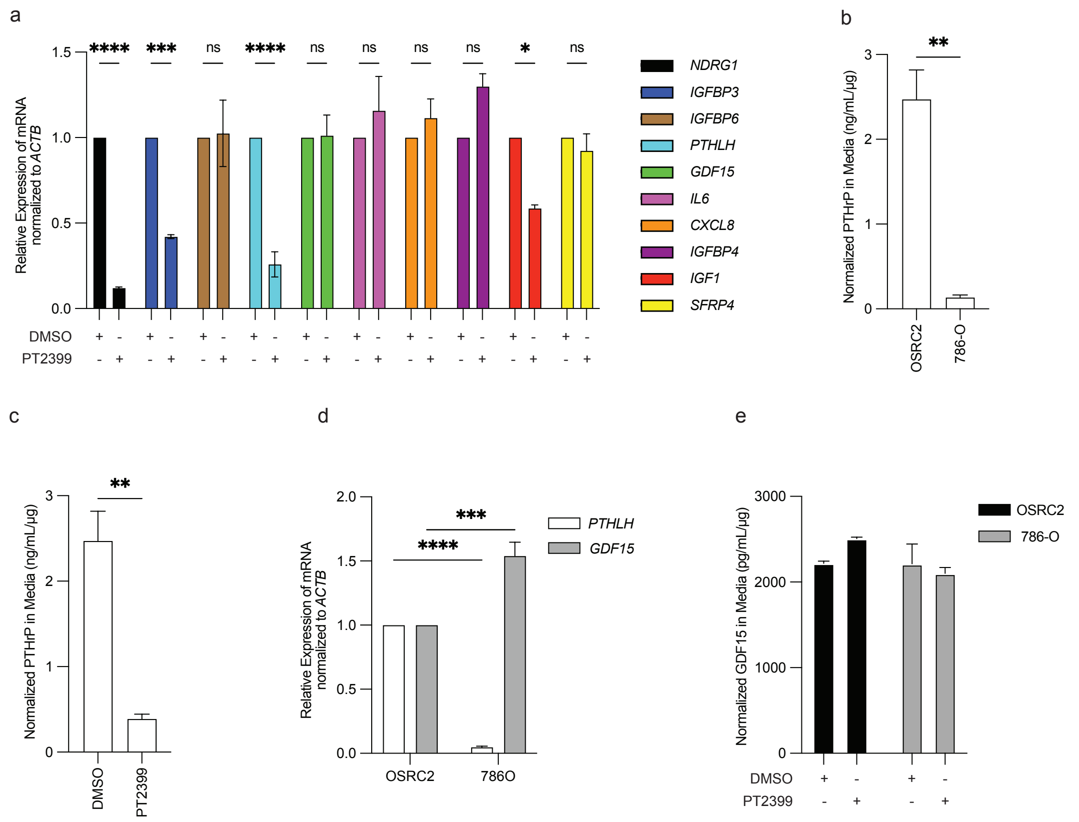


**Extended Data Figure 2. Validation of HIF2-responsive secreted proteins in OSRC-2 and 786-O cells.** (a) mRNA levels for selected HIF2-responsive genes (*NDRG1, IGFBP3, IGFBP6, PTHLH, GDF15, IL6, CXCL8, IGFBP4, IGF1, SFRP4*) in OSRC-2 cells treated with 2 mM PT2399 or DMSO for 48 hours. Each mRNA was normalized to *ACTB* and then to the DMSO value for that mRNA (n = 3). (b-c). PTHrP levels in media conditioned by OSRC-2 cells treated with 2 mM PT2399 or DMSO for 48 hours (b) or by OSRC-2 cells compared to 786-O cells (c) (n = 3). PTHrP levels were normalized to total cellular protein (ng/mL/μg). (d) Relative mRNA expression of *PTHLH* and *GDF15* in OSRC-2 and 786-O cells normalized to *ACTB* and then to the OSRC-2 value for that mRNA (n = 3). (e) GDF15 levels in media conditioned by OSRC-2 or 786-

O cells treated with 2 mM PT2399 or DMSO for 48 hours (n = 3). GDF15 levels were normalized to the total cellular protein (pg/mL/μg). Data are presented as mean ± SEM. Statistical significance was assessed using unpaired two-tailed t-tests or two-way ANOVA. P values are indicated as follows: $P \leq 0.05$ (*), $P \leq 0.005$ (**), $P \leq 0.005$ (***), and $P \leq 0.0005$ (****).

Extended Data Fig. 3

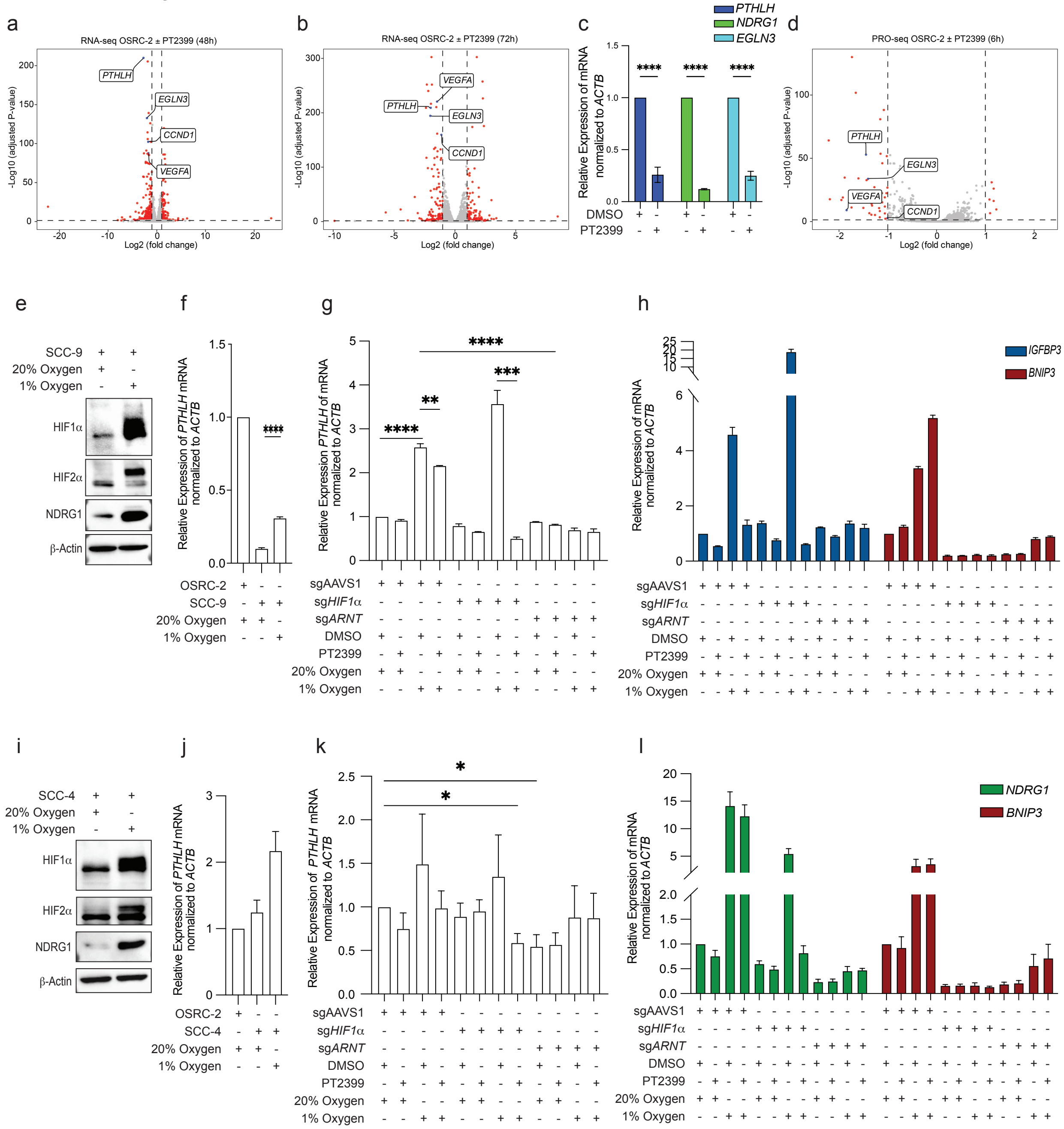


**Extended Data Figure 3. Extended analysis of *PTHLH* as a HIF2 target in OSRC-2 cells.** (a and b) Volcano plots of RNA-seq data comparing OSRC-2 cells treated with 2 mM PT2399 or DMSO for 48 hours (a) or for 72 hours (b). PTHLH and key HIF2 target genes, including *VEGFA, EGLN3*, and *CCND1*, are highlighted in blue. (c) Relative mRNA expression of *PTHLH, NDRG1,* and *EGLN3* in OSRC-2 cells treated with PT2399 or DMSO. Each mRNA was normalized to *ACTB* and then to the DMSO value for that mRNA (n = 3). (d) Volcano plot of PRO-seq data from OSRC-2 cells treated with PT2399 or DMSO for 6 hours, highlighting early transcriptional downregulation of *PTHLH, VEGFA, EGLN3,* and *CCND1*. (e,f and i,j) Immunoblot analysis (e and i) and qRT-PCR (f and j) of SCC-9 cells (e and f) or SCC-4 (i and j) incubated under 20% or 1% oxygen for 48 hours. *PTHLH* mRNA was normalized to *ACTB* and then to the OSRC-2 value (n = 3). (g,h and k,l) Relative mRNA expression of *PTHLH (g and k)*, or the HIF2 target gene *IGFBP3* and the HIF1$\alpha$ target gene *BNIP3 (h and l)*, in SCC-9 cells (g and h) or SCC-4 (k and l) edited with sgAAVS1, sg*HIF1*$\alpha$, or sg*ARNT*, treated with PT2399 or DMSO, and incubated under 20% or 1% oxygen levels for 48 hours. Each mRNA was normalized to *ACTB* and then to the sgAAVS1-DMSO-20% oxygen value for that mRNA (n = 3). Data are presented as mean ± SEM. Statistical significance was assessed using unpaired two-tailed t-tests and two-way ANOVA. P values are indicated as follows: *P ≤ 0.05* (**), P ≤ 0.005 (*****), P ≤ 0.005*** (***), and *P ≤ 0.0005* (****). For volcano plots, thresholds for significance were set at P ≤ 0.05, and n = 3. The adjusted p-value for *PTHLH* was zero but was set slightly above the smallest non-zero adjusted p-value of the other genes for graphical purposes.

### Extended Data Fig. 4

a

Amplicon sequencing sumary data of OSRC-2 sg*PTHLH*

| Alteration type | Position | Effect on coding sequence | Proportion (%) | Predicted impact |
|---|---|---|---|---|
| Deletion | Exon 6 | Deletion of 2 nucleotides | 53% | Early termination |
| Deletion | Exon 6 | Deletion of 1 nucleotides | 22% | Early termination |
| Deletion | Exon 6 | Framshift | 6% | Altered domain |
| Deletion | Exon 6 | Framshift | 6% | Truncated domain |

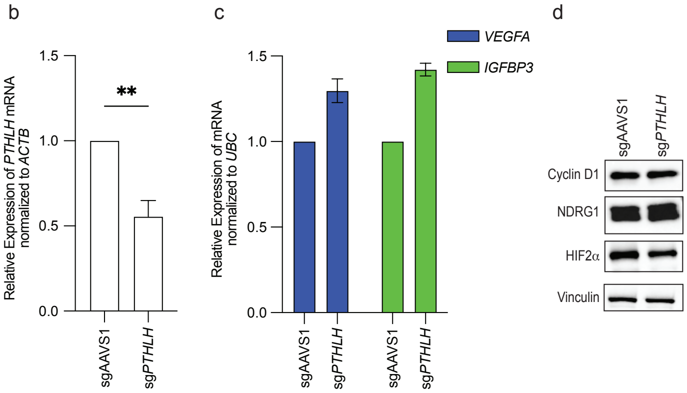


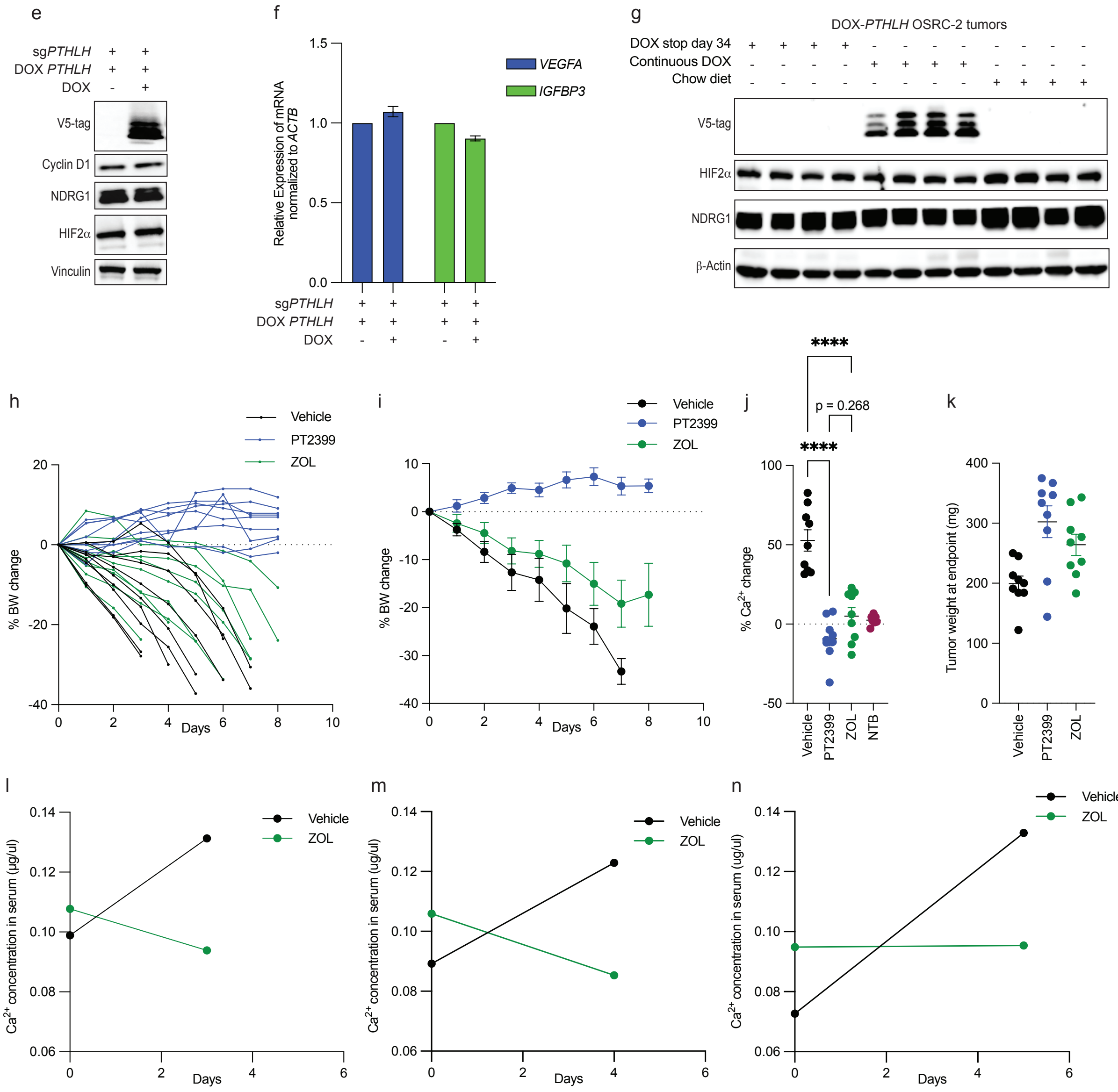

**Extended Data Figure 4. Characterization of OSRC-2 Cells after CRISPR-based editing with *PTHLH* sgRNA and Rescue with sgRNA-resistant *PTHLH* cDNA and Effects of Zoledronic Acid Treatment on Hypercalcemia and Cachexia** (a) Amplicon sequencing data summary for OSRC-2 cells subjected to CRISPR editing with a *PTHLH* sgRNA. (b-d). qRT-PCR (b and c) and immunoblot analysis (d) of OSRC-2 cells that underwent CRISPR with a *PTHLH* sgRNA or AAVS1 sgRNA. In (b and c) mRNA levels were normalized to *ACTB* and then to the *ACTB*-normalized (b) or *UBC*-normalized AAVS1 sgRNA cell value for that mRNA (n = 3). (e-g) immunoblot (e and g) and qRT-PCR (f) analysis of OSRC-2 sgPTHLH cells infected to express a DOX-inducible, sgRNA-resistant, *PTHLH* cDNA and grown in the presence or absence of doxycycline (DOX). In (f) mRNA levels were normalized to *ACTB* and then to the no DOX *ACTB*-normalized value for that mRNA (n = 3). (g) Immunoblot analysis of representative OSRC-2 tumor lysates at study endpoint in Fig. 4j. (h-i) Individual body weight (BW) trajectories (h) and group mean ± SEM for BW change over time (i) of mice treated with vehicle, PT2399, or Zoledronic acid (ZOL). (j) Percent change in serum calcium levels from baseline to endpoint in each treatment group vehicle, PT2399, or ZOL. NTB = non–tumor-bearing controls. (k) Tumor weights at endpoint for mice treated with vehicle, PT2399, or ZOL. (l–n) Early timepoint measurements of serum calcium from individual ZOL- and vehicle-treated mice that required early euthanasia due to cachexia. Data are presented as mean ± SEM. Statistical significance was assessed using unpaired two-tailed t-tests and two-way ANOVA test. P values are indicated as follows: *P ≤ 0.05* (*), *P ≤ 0.005* (**), *P ≤ 0.005* (***), and *P ≤ 0.0005* (****).

Extended Data Fig. 5

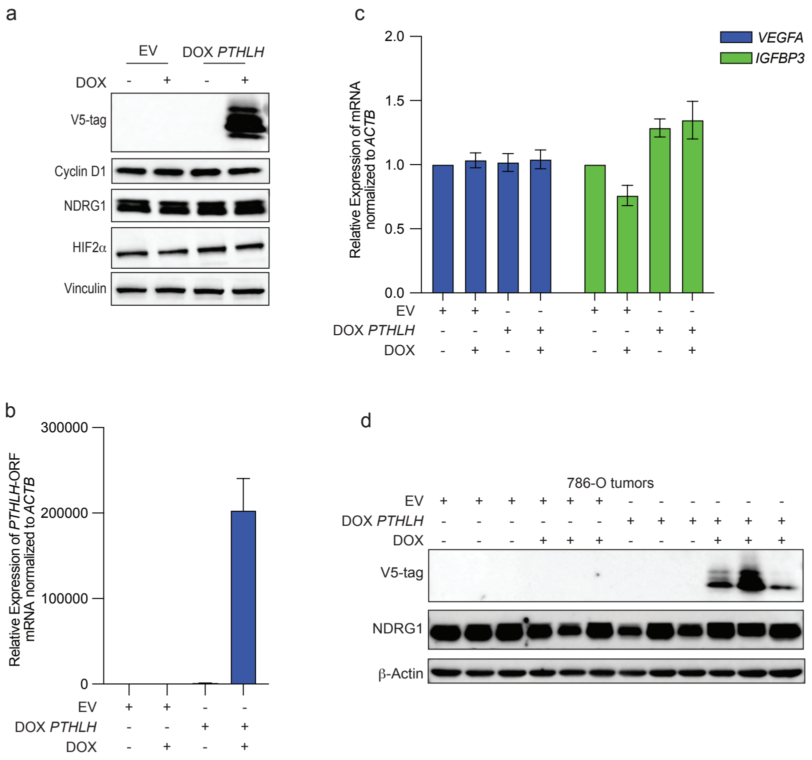


**Extended Data Figure 5. Extended analysis of *PTHLH* sufficiency for the induction of cachexia in 786-O xenografts.**

(a-c) Immunoblot (a) and qRT-PCR (c) analysis of 786-O cells infected with a doxycycline (DOX)-inducible *PTHLH* lentivirus or empty vector (EV). The *PTHLH* cDNA also encoded a 3' V5 epitope tag. Cells were grown in the presence of 1 mg/mL DOX for 3 days where indicated. In (b) qRT-PCR was performed with a primer targeting the V5 tag nucleotide sequence and mRNA values were normalized to *ACTB* then to the EV, no DOX, *ACTB*-normalized mRNA value (n = 3).  In (c) mRNA expression of *VEGFA* and *IGFBP3* was normalized to *ACTB* and then to the EV, no DOX, value for the corresponding mRNA (n = 3). (d) Immunoblot analysis of representative 786-O tumor lysates at study endpoint in Fig. 5e.

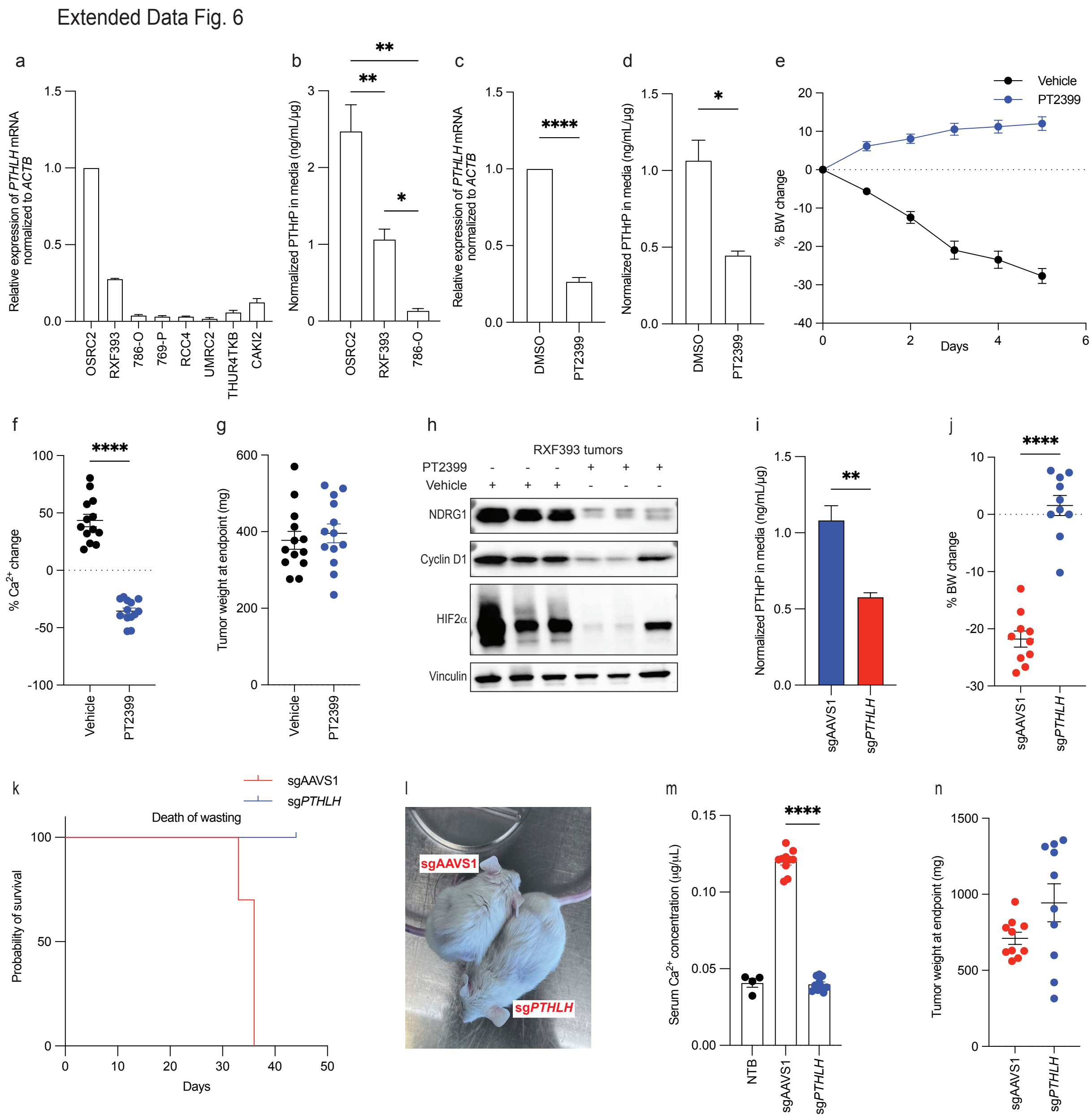


**Extended Data Figure 6. RXF393 induces cachexia via HIF2 and PTHrP.**

(a) *PTHLH* mRNA qRT-PCR across ccRCC cell lines. *PTHLH* mRNA levels were normalized to *ACTB* and then to the OSRC-2 cell *ACTB*-normalized value (n = 3). (b) PTHrP levels in media conditioned by OSRC-2, RXF393, or 786-O cells, normalized to total cellular protein (n = 3). (c-d) *PTHLH* mRNA qRT-PCR (c) and PTHrP ELISA (d) in OSRC-2 cells treated with 2m M PT2399 or DMSO for 48 hours. In (c) *PTHLH* mRNA levels were normalized to *ACTB* and then to the DMSO *ACTB*-normalized value (n = 3). In (b) PTHrP levels in the media was normalized to total cellular protein (n = 3). (e) Body weight trajectories of mice bearing RXF393 tumors treated with PT2399 (blue, n = 13) or vehicle (black, n = 13). (f-h) percent change in serum calcium levels relative to baseline pre-treatment levels (f), tumor weight at the study endpoint (g), immunoblot analysis of representative RXF393 tumor lysates at study endpoint in (h) of mice bearing RXF393 tumors treated with PT2399 or vehicle as in c). (i) PTHrP ELISA of 2 days conditioned media from RXF393 cells infected to express sg*PTHLH* or sgAAVS1, normalized to total cellular protein (n = 3). (j-k) Body weight change (j) and Kaplan-Meier survival curves (k) for mice bearing RXF393 tumors infected to express sg*PTHLH* (blue, n = 10) or sgAAVS1 (red, n = 10). (i-n) Representative mouse images (i), serum calcium levels (m) and tumor weight at study endpoint (n) of mice from (j). Statistical analysis for Kaplan-Meier was performed using the log-rank test (****$P \leq 0.0001$). Data are presented as mean ± SEM. Statistical significance was assessed using unpaired two-tailed t-tests or two-way ANOVA. P values are indicated as follows: $P \leq 0.05$ (*), $P \leq 0.005$ (**), $P \leq 0.005$ (***), and $P \leq 0.0005$ (****).

Extended Data Fig. 7

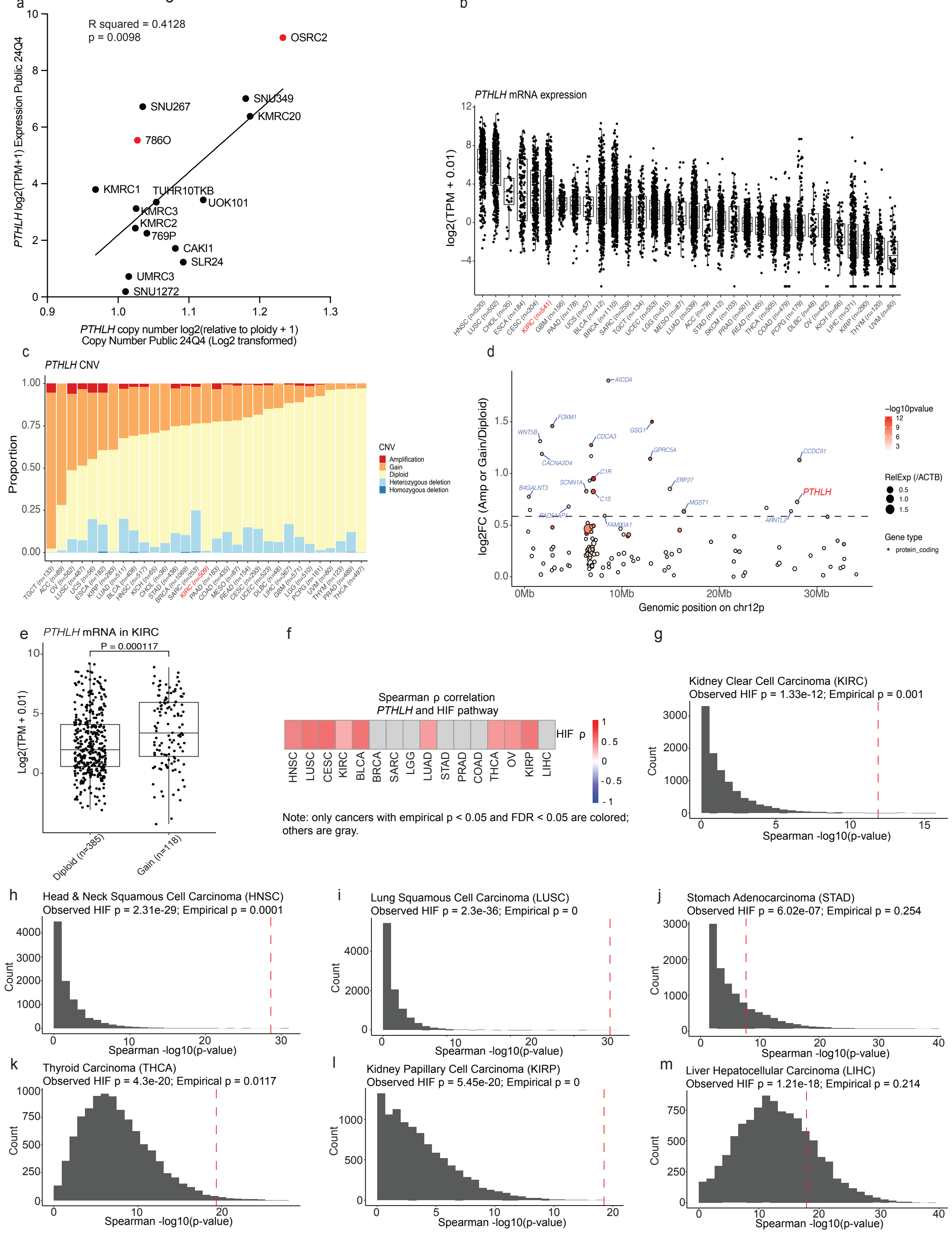

**Extended Data Figure 7. *PTHLH* is increased in KIRC, associated with copy number variation and gene expression in OSRC-2 and TCGA analysis, and correlates with a HIF transcriptional signature across multiple TCGA tumor types**

(a) Scatter plot showing the correlation between *PTHLH* gene copy number and mRNA expression in ccRCC cell lines. The x-axis shows *PTHLH* copy number (log2 relative to ploidy +1), and the y-axis shows *PTHLH* mRNA expression (log2 TPM +1). A linear regression curve was fitted using GraphPad Prism, with an $R^2$ value of 0.4128 and a p-value of 0.0098. OSRC-2 and 786-O are red labeled. Data extracted from Depmap.com 24Q4 dataset in Feb/09/2025. (b) Box plot of *PTHLH* mRNA expression across TCGA tumor types (log2 TPM+0.01), KIRC (Kidney Renal Clear Cell Carcinoma) is highlighted in red font. (c) Proportional bar plot showing *PTHLH* copy number variation (CNV) across TCGA tumor types. KIRC is highlighted in red font. (d) Scatter plot log2 fold change of mRNA expression of copy-gain/ copy-neutral samples for genes on chromosome 12p in TCGA KIRC samples, with *PTHLH* highlighted as a significantly amplified gene. Circle size represents relative expression (normalized to *ACTB*), and color intensity indicates -log10 p-value significance. The dashed line indicates fold change = 1.5. (e) Box plot showing *PTHLH* mRNA expression in TCGA KIRC samples with diploid (n = 385) or gain (n = 118) copy number states. Statistical significance (P = 0.000117) was assessed using unpaired Wilcoxon rank sum test. (f) Heatmap showing Spearman correlation coefficients (ρ) between *PTHLH* expression and the HIF transcriptional signature (*ADM*, *IRS2*, *EGLN3*, *EDN1*, *VEGFA*, *SLC2A1*, *TGFA*, *APOL1*, *CCND1*, *STC1*, *NDRG1*) across multiple TCGA cancer cohorts. Only associations meeting both statistical significance (FDR < 0.05) and empiric significance (empirical p-value < 0.05) are colored (red = positive correlation); non-significant associations are grey. (g-m) Examples of empiric p-value distributions for selected cancer types. For each cancer type, the observed correlation p-value between *PTHLH* and the HIF signature (red dashed line) was compared to the distribution of correlation p-values obtained from 10,000 random gene sets of identical size to the HIF signature. This approach addresses potential false positives and confirms that several cancer types including HNSC show robust, highly significant associations.

Extended Data Fig. 8

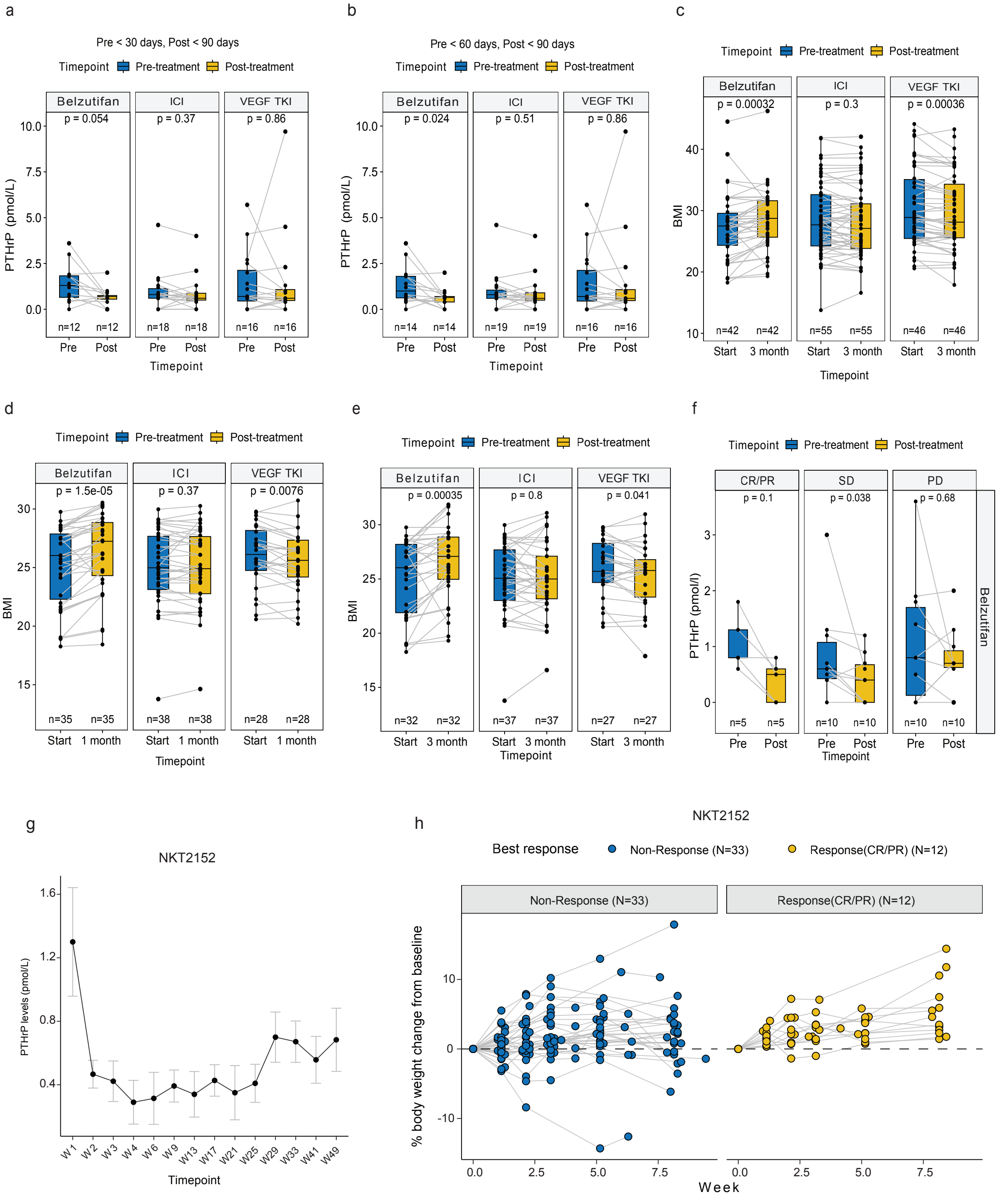

**Extended Data Figure 8. Additional analysis of hypercalcemia and cachexia treatments in RCC patients using HIF2 inhibitors, ICIs, and VEGF TKIs.**

(a and b) Plasma PTHrP levels in ccRCC patients treated with belzutifan (b: n = 12, c: n = 14), immune checkpoint inhibitors (ICIs; b: n = 18, c: n = 19), or VEGF tyrosine kinase inhibitors (TKIs; b: n = 16, b: c = 16) pre-treatment and post-treatment, stratified by different pre- and post-treatment time windows. (c) BMI in ccRCC patients treated with belzutifan (n = 42), ICIs (n = 55), or VEGF TKIs (n = 46), measured pre-treatment and after 3 months. (d-e) Changes in BMI for patients with pre-treatment BMIs <30 after being treated for 1 month (d) or 3 months (e) with belzutifan, ICI, or VEGF TKIs. (f) Plasma PTHrP levels in ccRCC patients treated with belzutifan, grouped by response category: CR/PR (complete/partial response, n = 5), SD (stable disease, n = 10), and PD (progressive disease, n = 10), measured pre-treatment and post-treatment. (g) Plasma PTHrP levels over time (up to Week 49) in ccRCC patients treated with NKT2152 (n = 14). Error bars represent standard error (SE). (h) Longitudinal changes in body weight (%) from baseline in patients treated with NKT2152, stratified by response status. Non-responders (n = 33; stable disease or progressive disease, blue) and responders (n = 12; partial or complete response, yellow). Data are presented as mean ± SEM. Statistical analysis and significance were assessed for (a-f) using the paired Wilcoxon rank-sum test.
