## Extended Data Table 1 for "HIF2-driven PTHrP Causes Cachexia and Hypercalcemia in Kidney Cancer: Treatment with HIF2 Inhibitors"

| | HIF2i (N=46) | ICI (N=56) | VEGF TKI (N=49) | Overall (N=151) | P-value |
|---|---|---|---|---|---|
| **Age at treatment start** | | | | | |
| Median IQR | 66.1 [60.0, 70.1] | 61.9 [57.4, 67.4] | 62.9 [55.9, 66.3] | 63.3 [57.2, 68.0] | 0.07 |
| **Sex** | | | | | |
| Female | 13 (28.3%) | 13 (23.2%) | 10 (20.4%) | 36 (23.8%) | 0.66 |
| Male | 33 (71.7%) | 43 (76.8%) | 39 (79.6%) | 115 (76.2%) | |
| **ECOG at treatment start** | | | | | |
| 0 | 27 (58.7%) | 40 (71.4%) | 27 (55.1%) | 94 (62.3%) | 0.31 |
| 1 | 14 (30.4%) | 13 (23.2%) | 16 (32.7%) | 43 (28.5%) | |
| 2 | 4 (8.7%) | 1 (1.8%) | 4 (8.2%) | 9 (6.0%) | |
| Missing | 1 (2.2%) | 2 (3.6%) | 2 (4.1%) | 5 (3.3%) | |
| **Best response** | | | | | |
| CR | 0 (0%) | 4 (7.1%) | 1 (2.0%) | 5 (3.3%) | 0.09 |
| PR | 9 (19.6%) | 18 (32.1%) | 17 (34.7%) | 44 (29.1%) | |
| SD | 19 (41.3%) | 20 (35.7%) | 23 (46.9%) | 62 (41.1%) | |
| PD | 15 (32.6%) | 11 (19.6%) | 7 (14.3%) | 33 (21.9%) | |
| Missing | 3 (6.5%) | 3 (5.4%) | 1 (2.0%) | 7 (4.6%) | |
| ***VHL* status** | | | | | |
| Wild-type | 4 (8.7%) | 5 (8.9%) | 6 (12.2%) | 15 (9.9%) | 0.918 |
| Mutated | 9 (19.6%) | 8 (14.3%) | 11 (22.4%) | 28 (18.5%) | |
| Missing | 33 (71.7%) | 43 (76.8%) | 32 (65.3%) | 108 (71.5%) | |
| **Stage** | | | | | |
| III | 0 (0%) | 3 (5.4%) | 1 (2.0%) | 4 (2.6%) | 0.23 |
| IV | 46 (100%) | 53 (94.6%) | 48 (98.0%) | 147 (97.4%) | |
| **Line of therapy** | | | | | |
| 1 | 0 (0%) | 47 (83.9%) | 37 (75.5%) | 84 (55.6%) | <0.001 |
| 2 | 5 (10.9%) | 4 (7.1%) | 9 (18.4%) | 18 (11.9%) | |
| 3 | 13 (28.3%) | 2 (3.6%) | 3 (6.1%) | 18 (11.9%) | |
| 4 | 13 (28.3%) | 2 (3.6%) | 0 (0%) | 15 (9.9%) | |
| 5+ | 15 (32.6%) | 1 (1.8%) | 0 (0%) | 16 (10.6%) | |
| **Corrected calcium at start of therapy (mg/dl)** | | | | | |
| Median IQR | 9.62 [9.40, 9.88] | 9.43 [9.12, 9.80] | 9.52 [9.28, 9.80] | 9.54 [9.27, 9.86] | 0.24 |
| **BMI at start of therapy (kg/m$^2$)** | | | | | |

| | HIF2i (N=46) | ICI (N=56) | VEGF TKI (N=49) | Overall (N=151) | P-value |
|---|---|---|---|---|---|
| Median IQR | 27.5 [24.2, 29.6] | 27.7 [24.1, 32.5] | 28.6 [25.5, 34.6] | 27.7 [24.9, 32.0] | 0.08 |

Table 1: Baseline characteristic of the patients with clear-cell renal cell carcinoma from the Dana-Farber cohort

CR: complete response, ECOG: Eastern Cooperative Oncology Group, HIF2i: HIF-2$\alpha$ inhibitors, ICI: immune checkpoint inhibitors, IQR: interquartile range, PD: progressive disease, PR: partial response, SD: stable disease, VEGF TKI: vascular endothelial growth factor tyrosine kinase inhibitor, *VHL*: Von-Hippel Lindau gene

Statistical comparisons between the 3 treatment groups were performed using the Chi-square test for categorical variables and the Kruskal-Wallis test for continuous variables.
