## Extended Data Table 2 for "HIF2-driven PTHrP Causes Cachexia and Hypercalcemia in Kidney Cancer: Treatment with HIF2 Inhibitors"

| | Low baseline PTHrP (≤ Q1) (N=12) | Mid baseline PTHrP (Q1- Q3) (N=22) | High baseline PTHrP (> Q3) (N=11) | Total Evaluable (N=45) |
|---|---|---|---|---|
| **Age at treatment start (years)** | | | | |
| Median [min-max] | 58 [47-78] | 67.5 [38-87] | 68 [53-74] | 64 [38-87] |
| **Sex** | | | | |
| Female | 2 (16.7%) | 9 (40.9%) | 4 (36.4%) | 15 (33.3%) |
| Male | 10 (83.3%) | 13 (59.1%) | 7 (63.6%) | 30 (66.7%) |
| **ECOG at treatment start** | | | | |
| 0 | 6 (50.0%) | 10 (45.5%) | 3 (27.3%) | 19 (42.2%) |
| 1 | 6 (50.0%) | 11 (50.0%) | 7 (63.6%) | 24 (53.3%) |
| 2 | 0 (0.0%) | 1 (4.5%) | 0 (0.0%) | 1 (2.2%) |
| Missing | 0 (0.0%) | 0 (0.0%) | 1 (9.1%) | 1 (2.2%) |
| **Best response** | | | | |
| CR | 0 (0.0%) | 0 (0.0%) | 0 (0.0%) | 0 (0.0%) |
| PR | 1 (8.3%) | 9 (40.9%) | 2 (18.2%) | 12 (26.7%) |
| SD | 6 (50.0%) | 10 (45.5%) | 4 (36.4%) | 20 (44.4%) |
| PD | 5 (41.7%) | 3 (13.6%) | 4 (36.4%) | 12 (26.7%) |
| Missing | 0 (0.0%) | 0 (0.0%) | 1 (9.1%) | 1 (2.2%) |
| **Stage** | | | | |
| IA | 1 (8.3%) | 0 (0.0%) | 0 (0.0%) | 1 (2.2%) |
| IB | 0 (0.0%) | 1 (4.5%) | 0 (0.0%) | 1 (2.2%) |
| IIA | 0 (0.0%) | 5 (22.7%) | 0 (0.0%) | 5 (11.1%) |
| IIB | 0 (0.0%) | 0 (0.0%) | 0 (0.0%) | 0 (0.0%) |
| IIIA | 6 (50.0%) | 6 (27.3%) | 4 (36.4%) | 16 (35.6%) |
| IIIB | 1 (8.3%) | 2 (9.1%) | 0 (0.0%) | 3 (6.7%) |
| IV | 2 (16.7%) | 4 (18.2%) | 7 (63.6%) | 13 (28.9%) |
| Other | 0 (0.0%) | 3 (13.6%) | 0 (0.0%) | 3 (6.7%) |
| Missing | 2 (16.7%) | 1 (4.5%) | 0 (0.0%) | 3 (6.7%) |
| **Line of therapy** | | | | |
| 1 | 3 (25.0%) | 3 (13.6%) | 0 (0.0%) | 6 (13.3%) |
| 2 | 3 (25.0%) | 3 (13.6%) | 4 (36.4%) | 10 (22.2%) |
| 3 | 2 (16.7%) | 8 (36.4%) | 3 (27.3%) | 13 (28.9%) |

| | Low baseline PTHrP (≤ Q1) (N=12) | Mid baseline PTHrP (Q1- Q3) (N=22) | High baseline PTHrP (> Q3) (N=11) | Total Evaluable (N=45) |
|---|---|---|---|---|
| 4 | 2 (16.7%) | 2 (9.1%) | 2 (18.2%) | 6 (13.3%) |
| 5 | 0 (0.0%) | 2 (9.1%) | 1 (9.1%) | 3 (6.7%) |
| 6+ | 2 (16.7%) | 4 (18.2%) | 1 (9.1%) | 7 (15.6%) |
| **Corrected calcium at start of therapy (mmol/L)** | | | | |
| Median [min-max] | 2.29 [2.12-2.62] | 2.38 [2.07-2.62] | 2.45 [2.26-3.45] | 2.40 [2.07-3.45] |
| **Body weight at start of therapy (kg)** | | | | |
| Median [min-max] | 97.6 [61.6-127] | 75.1 [49.6-117] | 84.6 [60.3-94.3] | 81.8 [49.6-127] |

Table 2: Baseline characteristic of the patients with clear-cell renal cell carcinoma from the Nikang cohort

CR: complete response, ECOG: Eastern Cooperative Oncology Group, PD: progressive disease, PR: partial response, PTHrP: parathyroid hormone-related protein, Q1: first quartile, Q2: second quartile, Q3: third quartile, SD: stable disease.
